# Structural basis of the nucleotide incorporation cycle of bacterial DNA polymerase III

**DOI:** 10.64898/2026.06.29.735100

**Authors:** Zhi-Qiang Xu, Slobodan Jergic, Qiang Zhu, Haibo Yu, Jacob Lewis, Simon Brown, James Bouwer, Harshad Ghodke, Peter J. Lewis, Gökhan Tolun, Aaron Oakley, Nicholas E. Dixon

**Affiliations:** Molecular Horizons and School of Science, University of Wollongong, New South Wales 2522, Australia; ARC Centre of Excellence in Quantum Biotechnology, University of Wollongong, New South Wales 2522, Australia; Hunter Biological Solutions, 1/82-84 Beaumont Street, Hamilton, New South Wales 2303, Australia; ARC Industrial Transformation Training Centre for Cryo-electron Microscopy of Membrane Proteins, University of Wollongong, New South Wales 2522, Australia

## Abstract

Bacterial DNA polymerase III (PolIII) is the primary replicative polymerase responsible for faithful genome duplication, yet high-resolution structural snapshots of the enzyme during DNA synthesis and proofreading have remained elusive. Here we determine cryo-EM structures of the *Escherichia coli* PolIII core–clamp–DNA binary, ternary and proofreading complexes at 2.4–2.7 Å resolution, revealing the structural basis of nucleotide selection, incorporation and proofreading. Binding of an incoming nucleotide expels the single-stranded template overhang from the polymerase central channel and induces a large rotation of the polymerase index finger, which closes around the incoming nucleotide to form a compact nascent base-pair-binding pocket. A conserved tyrosine on the finger stacks against the incoming nucleotide to impose stringent steric constraints that promote nucleotide selection, whereas a conserved histidine acts as a steric gate to exclude ribonucleotides. The proofreading structures capture key intermediates that reveal how PolIII transfers a mismatched primer terminus to the 3′–5′ exonuclease active site and excises the misincorporated nucleotide while avoiding unnecessary degradation of correctly paired DNA. Together, these findings define the molecular mechanism by which the bacterial replicative polymerase couples rapid and processive DNA synthesis with exceptional replication fidelity.

## INTRODUCTION

DNA polymerase III (PolIII) replicates the bacterial genome in a fast, highly processive manner with high fidelity (accuracy). The *Escherichia coli* PolIII holoenzyme (HE) can continuously synthesize tens of kilo-base pairs at a speed of ∼1000 nucleotides (nt) per second^1^ with an error rate of ∼1 per 10^6^ nucleotide incorporations^2^. The structural dynamics of PolIII that enables this feat remains largely unknown due to the lack of high-resolution structures of PolIII in complex with a primer-template (p/t) DNA and an incoming nucleotide.

PolIII, a C-family DNA polymerase, is structurally and evolutionarily distinct from the viral, eukaryotic, and archaeal replicative polymerases that belong to A, B and D families^3–7^. An *E. coli* PolIII HE contains two to three DNA polymerase cores (αεθ) that synthesize DNA, the β_2_ DNA sliding clamps that tether polymerase cores to DNA, thus substantially increasing their speed and processivity, and a seven-subunit clamp loader complex (CLC, τ_n_γ_3-n_δδ′ψχ; n=2−3) that opens and loads β_2_ clamps onto p/t DNA^1,8–10^. PolIII core comprises a polymerase subunit α, a 3′−5′ exonuclease ε and a θ that stabilizes ε. Both α and ε contain a clamp-binding motif (CBM) that binds to one of the two identical CBM-binding pockets on β_2_^1,11,12^. Both τ and γ of the CLC are products of the *dnaX* gene. τ is a full-length product while γ is shorter due to ribosome translational frameshift. The extra C-terminal Domain IV of τ interacts with the DnaB DNA helicase that unwinds double-stranded (ds) DNA and the 16 kD Domain V (τ_C16_)^13^ binds tightly to the C-terminal τ-binding domain of α (TBD), through which PolIII HE and the DNA helicase can work as a highly orchestrated molecular machine. The structure of α resembles a cupped right hand, comprised of the N-terminal PHP (polymerase and histidinol phosphatase) domain, palm, thumb, fingers, Linker, OB (oligonucleotide/oligosaccharide binding) domain and TBD^5,6^ (Supplementary Fig. 1A,B). The finger domain of α is much larger than those of the A- and B-family polymerases that is further divided into the index, middle, ring, and little fingers. The ε subunit contains a globular exonuclease domain (2−186, ε186) and a poorly conserved 63-residue C-terminal segment, the last ∼40 residues of which (ε_CTD_) binds tightly to the PHP^14^.

So far, high-resolution structural and mechanistic insights into how the replicase engages DNA, selects and incorporates nucleotides, and corrects replication errors remain elusive, although several structures of bacterial replicative DNA polymerases have been reported, including apo α from *E. coli* (2HNH)^5^ and *Thermus aquaticus* (2HPI)^6^, the low-resolution *E. coli* PolIII binary (5FKV, 8 Å)^15^ and proofreading complexes (5M1S, 6.7 Å)^16^, and the *T. aquaticus* ternary complex (3E0D, 4.6 Å)^17^ (Supplementary Fig. 1B−D). A structure of the Gram-positive PolC ternary complex (3F2B, 2.4 Å) is also available^18^ (Supplementary Fig. 1E). However, PolC diverged from PolIIIα more than a billion years ago, sharing little sequence identity with different functional modules and domain organization (Supplementary Fig. 1A).

Here, we sought to determine high-resolution structures of the *E. coli* PolIII core−clamp−DNA complexes. We systematically characterized the α−DNA interactions using surface plasmon resonance (SPR) analysis, engineered both α and ε to stabilize the complexes for cryo-EM analysis, and determined structures of the PolIII binary, ternary and proofreading complexes at 2.4−2.7 Å. Together with biochemical analyses and molecular dynamics (MD) simulations, these structures define the structural basis of nucleotide selection, incorporation and proofreading by PolIII, revealing the molecular mechanism underlying a dNTP incorporation cycle by the bacterial replicative DNA polymerase at near-atomic resolution.

## RESULTS

### Stabilization of the *E. coli* PolIII αεθ·β_2_·DNA complex for cryo-EM

To stabilize the αεθ·β_2_·DNA complexes for structure determination, we first tested how the α^E612K^ variant that was known to bind to DNA more tightly than wild-type α^11,19–21^ binds to p/t DNAs with single-stranded (ss) template overhangs ranging from 3 to 15 nucleotides (nt) by SPR. We found that α^E612K^ bound increasingly more tightly to p/t DNAs with longer overhangs. The binding affinities (*K*_D1_) increased ∼16 fold as the overhang lengthened from 3 to 15 nt in the absence of incoming nucleotide (dGTP) (*i.e*., 3800 nM *vs.* 240 nM) (Table 1, Supplementary Figs. 2−4, Supplementary Data 1 and 2). The presence of dGTP further strengthened α^E612K^–DNA interaction by up to 17-fold (*i.e*., *K*_D_^app^/*K*_D1_ value for p/t DNA with a 15-nt overhang). We then replaced the CBMs of both α^E612K^ and ε^D12A/E14A^ that has no exonuclease activity^11^ with stronger binding motifs, *i.e*., QLDLF for α^E612K^ (α^EL^) and QLSLPL for ε^D12A/E14A^ (ε^DL^), to strengthen binding of αεθ to β_2_. These mutations are not expected to modify the mechanism of nucleotide selection and incorporation. Indeed, α^EL^ was found to be more active than wild-type α in DNA synthesis (Supplementary Fig. 5). The resultant α^EL^ε^DL^θ·β_2_·DNA complex was stable enough to be isolable on size-exclusion chromatography in the presence of dGTP (Supplementary Fig. 6). Using the α^EL^ε^DL^θ complex, the structures of the PolIII binary (αεθ·β_2_·DNA), ternary (αεθ·β_2_·DNA·dGTP), and binary-like pre-proofreading intermediate and proofreading (αεθ·β_2_·DNA^mismatch^) complexes were subsequently determined by single-particle cryo-EM.

**Table 1.** Equilibrium binding parameters of α^E612K^ and wild-type α binding to p/t DNAs with ss template overhang of various lengths in the absence and presence of dGTP, based on SPR data in Supplementary Figures 2, 3 and 4. Dissociation constants for the ^bio^p/t DNA−α interactions (*K*_D1_) and apparent dissociation constant (*K*_D_^app^) for the ^bio^p/t DNA−α interactions in the presence of 0.5 mM dGTP were determined by fitting responses at equilibrium in Supplementary Figures 2, 3 and 4 to Equation 1 (see Methods) and Equation 8 (Supplementary Data 1), respectively. The apparent dissociation constant *K*_D2_^app^ that broadly reports on the nucleotide affinity was calculated from Equation 10 (Supplementary Data 1). The errors are standard errors of the fits. 0−2 nt indicates a total of 2 nt on the template overhang in the presence of incoming nucleotide where 0 nt denotes templating nucleotide, or 3 nt in the absence of incoming nucleotide. Thorough description of SPR analysis is included in Supplementary Data 1 and 2.

| Reaction | ss DNA | p/t DNA<br>0–2 nt | p/t DNA<br>0–4 nt | p/t DNA<br>0–5 nt | p/t DNA<br>0–6 nt | p/t DNA<br>0–7 nt | p/t DNA<br>0–9 nt | p/t DNA<br>0–14 nt |
| --- | --- | --- | --- | --- | --- | --- | --- | --- |
| **$K_{D1}$ (nM)**<br>$\alpha^{E612K}$-DNA | 1200<br>$\pm 130$ | 3800<br>$\pm 400$ | 1800<br>$\pm 200$ | 1300<br>$\pm 100$ | 630<br>$\pm 50$ | 430<br>$\pm 30$ | 280<br>$\pm 30$ | 240<br>$\pm 20$ |
| **$K_{D}^{app}$ (nM)**<br>$\alpha^{E612K}$-DNA + dGTP | 890<br>$\pm 80$ | 3500<br>$\pm 400$ | 890<br>$\pm 50$ | 370<br>$\pm 20$ | 68<br>$\pm 6$ | 43<br>$\pm 4$ | 20<br>$\pm 4$ | 14<br>$\pm 2$ |
| **Calculated $K_{D2}^{app}$ (μM)**<br>$\alpha^{E612K}$.DNA-dGTP | – | > 1535 | 500<br>$\pm 200$ | 190<br>$\pm 50$ | 61<br>$\pm 12$ | 56<br>$\pm 10$ | 42<br>$\pm 14$ | 32<br>$\pm 8$ |
| **$K_{D1}$ (nM)**<br>wt α-DNA | – | – | – | – | – | – | 3800<br>$\pm 400$ | 3100<br>$\pm 200$ |
| **$K_{D}^{app}$ (nM)**<br>wt α-DNA | – | – | – | – | – | – | 1110<br>$\pm 60$ | 820<br>$\pm 50$ |
| **Calculated $K_{D2}^{app}$ (μM)**<br>wt $\alpha$.DNA-dGTP | – | – | – | – | – | – | 210<br>$\pm 50$ | 180<br>$\pm 30$ |

### The template overhang resides in the central channel of α in “ground-state” binary complex

First, we solved the structures of the *E. coli* αεθ·β_2_·DNA complex bound to the C-terminal 32-kD fragment of τ that contains both Domains IV and V (τ_C32_)^22^ and a p/t DNA containing a 23-bp region and a (dT)_10_ ss template overhang in the absence of incoming nucleotide (binary complex, PoIIII^BC^). Three cryo-EM reconstructions at 2.5 Å revealed well-resolved N-terminal domains of α (NTD, 1–968), containing the PHP, thumb, palm, fingers and Linker, bound to a p/t DNA and β_2_ (Supplementary Table 1, Supplementary Fig. 7, Figure 1A). OB is less-well resolved, while TBD and its binding partner τ_C32_ could not be detected, indicating a flexible connection with the rest of α. While the C-terminal PHP-binding peptide of ε (ε_CTD_)^14^ is well-resolved, densities of the nuclease domain of ε (ε186)^23^ and θ are much weaker but still allow docking of the known εθ structure (2XY8)^24^. The three structures are similar (αs superimposed with RMSDs of 0.480−0.722 over 1080 Cα pairs) with β_2_ tilted and rotated and εθ shifted relative to α, which were assigned as PoIIII^BC1−3^ based on the α−β_2_ angle (Supplementary Fig. 7A).

**Figure 1.**
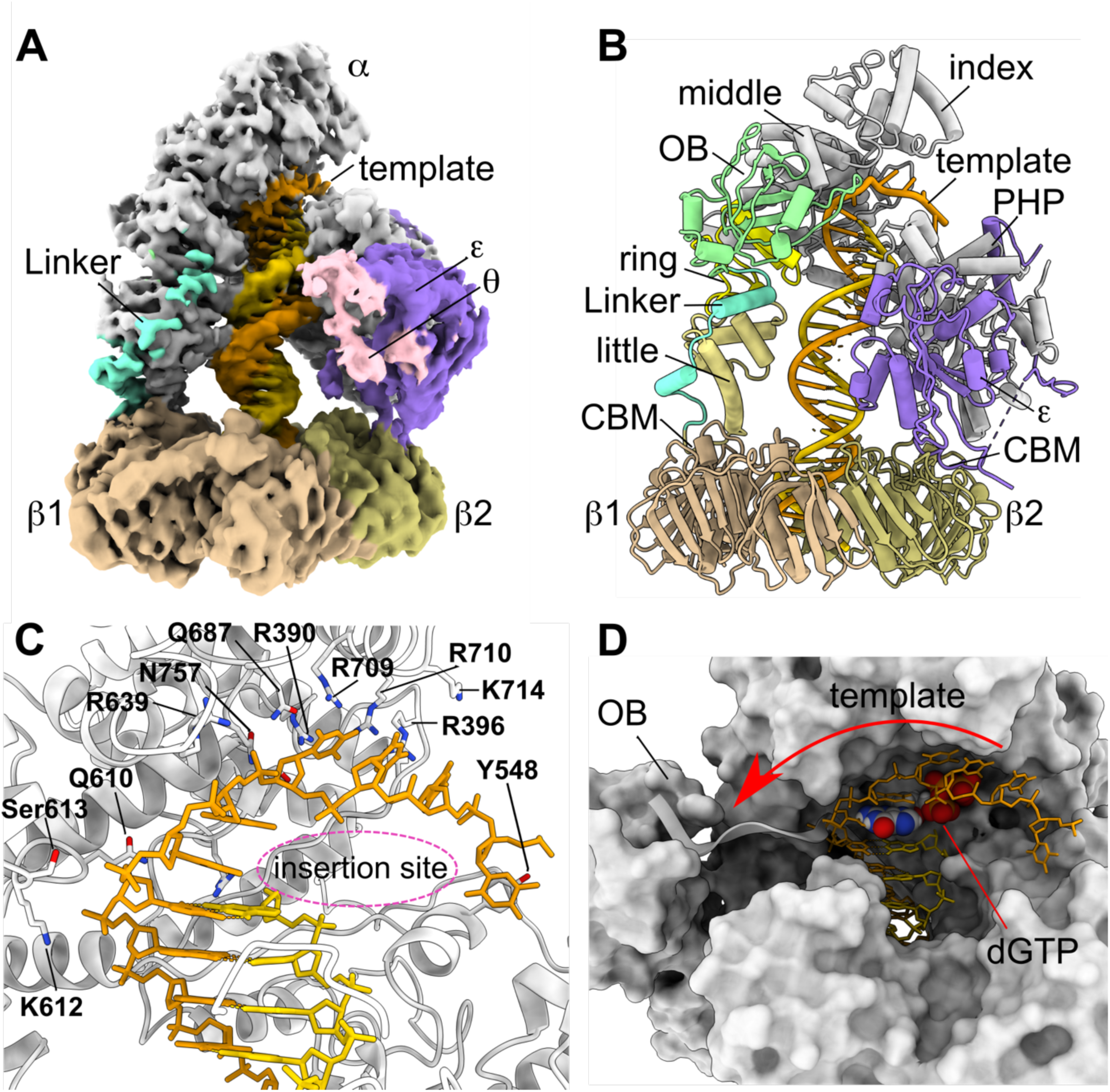
Structures of the *E. coli* PolIII binary complexes (PolIII^BC^). (**A**), Cryo-EM map of the *E. coli* PolIII^BC1^ complex with visualized densities corresponding to αNTD (grey), Linker connecting the little finger and OB (aquamarine), ε (medium purple), θ (pink), β_2_ (dark khaki and tan), and primer (gold) and template strands (orange), respectively. Density of OB is weak that can only be seen at low density threshold. (**B**), Structure of PolIII^BC1^, showing that the Linker of α (aquamarine) binds to both little (khaki) and ring (yellow) fingers to place OB (light green) near the polymerase active site. (**C**), The single-stranded template overhang of PolIII^BC^ (orange) resides in the central channel of α. Several positively charged and polar residues (sticks and labelled) on the surface of the central channel may interact with and stabilize the template overhang. (**D**), The template overhang in the central channel of PolIII^BC^ (orange sticks) would potentially clash with incoming nucleotide (spheres); and there is no steric hindrance in α^BC^ preventing the overhang from swinging to the side path (red arrow and white ribbon).

The structures of PolIII^BC^ provide substantially more detailed information than the 8 Å binary structure (*Eco*PoIIII^BC^, 5FKV)^15^. Notably, the template overhang occupies the central channel of α, precluding formation of a nascent base pair-binding pocket. However, no steric hindrance appears to prevent the overhang from swinging outward (Figure 1B−D). Six unpaired nucleotides are seen reaching towards the PHP near where ε binds (Figure 1C), stabilised by positively charged and polar residues from the palm, index and middle fingers, such as Arg390, Arg396, Arg639, Arg709 and Arg710. The base of the last visible nucleotide may stack on the benzene ring of Tyr548. Unlike *Eco*PoIII^BC^, the structurally rigid Linker positions the OB near the polymerase active site rather than away from it (Supplementary Fig. 1D). The Linker folds into two α-helices, the first tracks along the little finger and the second binds at the interface between the little and ring fingers, bundling them together.

α−DNA interactions are mostly restricted to the front part of α, mainly through palm and thumb that hold DNA inside α (Supplementary Fig. 8). The ds DNA passes through the β_2_ clamp nearly perpendicular to the ring (∼81°) and does not interact with β_2_ extensively, interacting mostly via a few positively charged residues, including Arg24, Arg73 and Arg205. Interactions among protein subunits are not extensive either, except the αPHP–ε_CTD_ contact^14^, and the α–β_2_ and ε–β_2_ interactions via CBMs. εθ is closer to α in PolIII^BC1^ than PolIII^BC2,3^ with the globular domain of ε contacting thumb while θ might also contact OB.

### α reorganizes extensively to bind DNA

Comparing α^BC^ with α^apo^ (2HNH), conformational changes of α induced by DNA binding are pronounced. Most domains of α undergo significant movements and/or conformational changes. Specifically, fingers move closer to DNA; the PHP, thumb, and part of the palm that blocks DNA binding in α^apo^ rotate clockwise to accommodate DNA binding (Figure 2A,B, Movie 1). On the most conserved palm domain, while the N-terminal region (271-393) of the large palm sub-domain (palm^L^, 271–432) is mainly unchanged, the remaining part carrying catalytic Asp401 and Asp403 residues undergoes significant conformational changes (Figure 2B). The small palm sub-domain (palm^S^, 511–560) carrying the third catalytic carboxylate Asp555 rotates as a rigid body along with the PHP and thumb. The rotation reduces the distance between Cα atoms of Asp403 and Asp555 from 7.2 Å in α^apo^ to 4.7 Å in α^BC^, restoring the otherwise distorted β-NT domain^6,17^.

**Figure 2.**
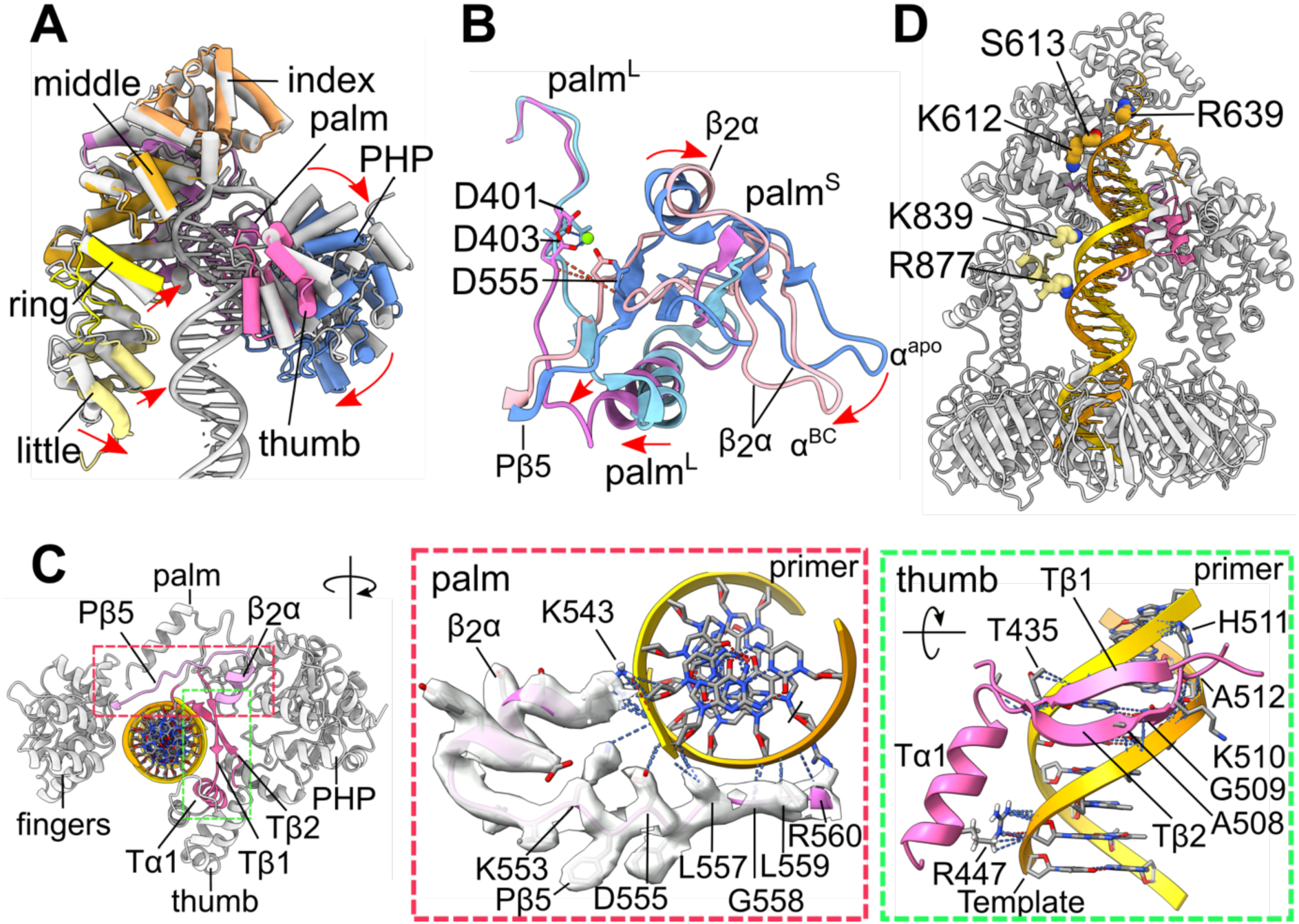
Conformational changes of α induced by DNA binding. (**A**), Superimposition of α^BC^ and α^apo^ (2HNH), showing significant conformational changes on DNA binding. Arrows indicate translational or rotational movements of individual domains upon DNA binding to avoid steric clashes and increase interactions with DNA. PHP, palm, thumb, and index, middle, ring and little fingers of α^BC^ are coloured cornflower blue, orchid, hot pink, sandy brown, goldenrod, and yellow, respectively. α^apo^ and DNA are coloured grey. (**B**), Conformational changes of the palm domain. Rotation of palm^S^ reduces the distances (orange dash lines) between Cα atoms of Asp403 and Asp555 from 7.2 Å in α^apo^ to 4.7 Å in α^BC^, restoring the otherwise distorted β-NT domain in α^apo^. The large palm sub-domain (palm^L^) of α^BC^ and α^apo^ are coloured orchid and sky blue, respectively; and the small palm sub-domain (palm^S^) of α^BC^ and α^apo^ are coloured light pink and cornflower blue, respectively. (**C**), Interactions of the palm and thumb with DNA. Left, top view of the DNA-bound α with the index and middle fingers and part of the palm omitted. Middle, the C-terminal β strand of the palm (Pβ5) and the conserved β_2_α structural motif (pink ribbons with electron densities as surface) that are in steric conflict with DNA binding in α^apo^ move sideways to allow DNA binding, whereby they interact extensively with the backbones of both primer (gold) and template strands (orange). Only the helix of the β_2_α motif is shown. α−DNA interactions are indicated by dashed lines. Right, interactions of the anti-parallel β strands of the thumb with DNA. (**D**), Interactions between fingers and DNA. Residues on the middle and little fingers that directly contact DNA are shown as spheres and coloured goldenrod and khaki, respectively.

The palm and thumb are most extensively involved in DNA interactions in α (Supplementary Fig. 8). The C-terminal β-strand of the palm (Pβ5, Phe554–Arg560) that would clash with DNA binding in α^apo^ moves sideways to accommodate DNA binding, whereby it interacts with the backbones of the minor groove of the DNA (Figure 2C, Supplementary Movie 1). Gly558, Leu559 and R560 contacts the −3 nt of the template strand, while Leu557, Asp555 and Lys553 interact with −1 nt of the primer strand. Lys543 on the conserved β_2_α structural motif inserted between Pβ4 and Pβ5^5^ interacts with −1 and −2 nt of the primer strand, forming a hydrogen bond with phosphate oxygen of −1 nt.

The highly conserved antiparallel β-strands of the thumb (Tβ1 and Tβ2)^18^ also move sideways and interact with the minor groove of DNA. Tβ1 (Gln429–Ala438) interacts mainly with the backbone of −3 and −4 nt of the primer and Tβ2 (Gly503–Gly513) interacts with −4 to −6 nt of the template (Figure 2C). Conserved residue His511 interacts with −2 nt of the primer, consistent with a lack of polymerase activity in the K510A-H511A double mutant^25^. The helix following Tβ1 (Tα1, Lys439−Pro452) contacts backbones of both DNA strands, with Arg447 hydrogen bonding to −8 and −9 nt of the template.

Interactions between DNA and the fingers are less extensive and mainly through the backbone of the DNA. Loop Phe638–Ser645 between the index and middle fingers interacts with the template overhang while loop Phe609–Ser613 of the middle finger contacts the template near the ss-ds DNA junction (Figure 2D, Supplementary Fig. 8). Lys612 interacts with −2 nt that explains why α^E612K^ binds more tightly to p/t DNA and was more active than wild-type α (Table 1, Supplementary Figure 5). Loop Ile838–Gly844 of the little finger contacts the primer, while helix Asn875–Met884 interact with the template (Figure 2D).

### The template overhang swings to side path upon incoming nucleotide binding

Next, we solved the structures of the αεθ·τ_C32_·β_2_·DNA·dGTP ternary complex (PoIIII^TC^) using a p/t DNA with a 2′,3′-dideoxycytidine monophosphate (ddCMP) at the 3′-end of the primer to prevent strand extension. The overall structures of the four PoIIII^TC^ complexes (Supplementary Table 2, Supplementary Fig. 9) are similar to PoIIII^BC^. The αNTD, Linker and DNA are again well-resolved, while densities of OB and εθ are weaker, and the TBD and τ_C32_ are missing (Figure 3A−C). However, the OBs are more clearly defined than in PoIIII^BC^, likely due to direct contacts with the template overhang. We also solved the structures of PolIII^TC^ using a full τ_3_δδ′ψχ CLC instead of τ_C32_ with the same results (data not shown).

**Figure 3.**
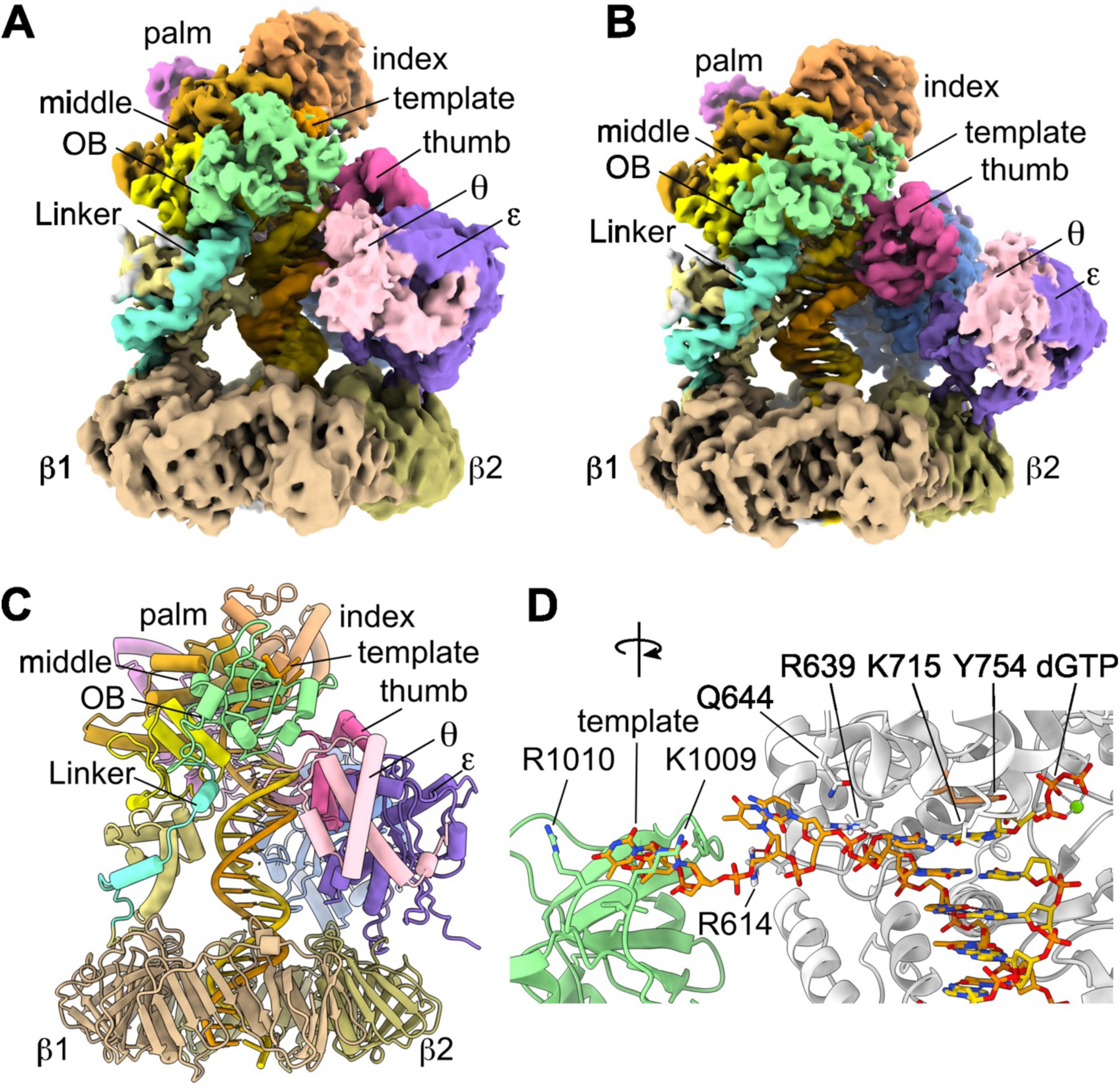
Structures of the *E. coli* PolIII ternary complexes (PolIII^TC^). (**A**), Cryo-EM map of PolIII^TC2^. (**B**), Cryo-EM map of PolIII^TC4^. PolIII^TC4^ appears more compact than PolIII^TC2^. Densities corresponding to different domains of α: the PHP, palm, thumb, and index, middle, ring and little fingers, Linker and OB are coloured cornflower blue, orchid, hot pink, sandy brown, goldenrod, yellow, khaki, aquamarine and light green, respectively. Density corresponding to ε is coloured medium purple, θ pink, the β subunits dark khaki and tan, primer strand gold and template strand orange. (**C**), Structure of PolIII^TC2^. (**D**), The ss template overhang is sharply kinked around the templating nucleotide and passes through a surface groove of OB, whereby it may interact with Ly1009 and Arg1010 on OB. The overhang interacts with residues Arg614, Gly615, Arg639, Gln644, and Lys715. Arg639 hydrogen bonds (red dashed lines) to both 0 and +1 nt of the template strand and Arg614 to +2 nt.

The four PoIIII^TC^ structures are similar with subtle movements on the α subunit relative to DNA and other proteins. Aligning on α, the ds DNA tilts or bends at slightly different angles, while β_2_ tilts and rotates to remain nearly perpendicular to the DNA (Supplementary Movie 2). These movements make these complexes looking increasingly more compact.

Although the α−ds DNA interactions in PoIIII^TC^ closely resemble those in PoIIII^BC^ (Supplementary Fig. 10), a sharp kink in the template strand near the templating nucleotide redirects the overhang into the DNA side path, as observed in the *T. aquaticus* ternary complex (*Taq*PolIII^TC^, 3E0D)^17^ (Figure 3D). Arg639, which hydrogen bonds with the template strand at the ds-ss DNA junction, may chaperon the overhang during its outward movement. Four unpaired nucleotides (+1 to +4) ahead of the templating nucleotide are visible (Figure 3D). The overhang does not interact with α extensively and is just long enough to reach the surface groove of OB flanked by moderately conserved basic residues, Lys1009 and Arg1010^5^. This is consistent with the proposed role of OB in guiding a ss template strand into the polymerase active site^17^. No density for nucleotides beyond +4 nt is clearly identified, suggesting that DNA beyond this point is not rigid enough to be resolved. In agreement with this, our SPR data show that binding affinity (*K*_D_^app^) of α^E612K^ to p/t DNA with a 4-nt overhang (“0−4 nt” p/t DNA) is ∼4-fold stronger than 2-nt in the presence of cognate nucleotide (Table 1 and Supplementary Data 1 and 2). An overhang longer than 4 nt still contributed to α−DNA binding with the apparent binding affinity increasing more than 60-fold when the overhang increased from 4 to 14 nt. These data, together with the fast kinetics seen in sensorgrams (Supplementary Fig. 3), point to the dynamic and transactional nature of the α−ss DNA interactions that may allow ss DNA to slide on the surface of α (OB), rather than forming a static complex.

The position of εθ shifts substantially in different complexes (Supplementary Movie 2), consistent with the notion that ε is flexibly tethered to β_2_^14^. However, the nuclease active sites of ε always faces DNA and are accessible for proofreading.

### The index finger of α closes around dGTP to form a compact nascent base pair-binding pocket

Although the overall structure of α^TC^ is similar to that of α^BC^, the index finger undergoes a large rotation to close around the dGTP (Figure 4A, Movie 2). We refer to the index finger conformation in α^TC^ as the “closed” state and that in α^BC^ as the “open” state.

**Figure 4.**
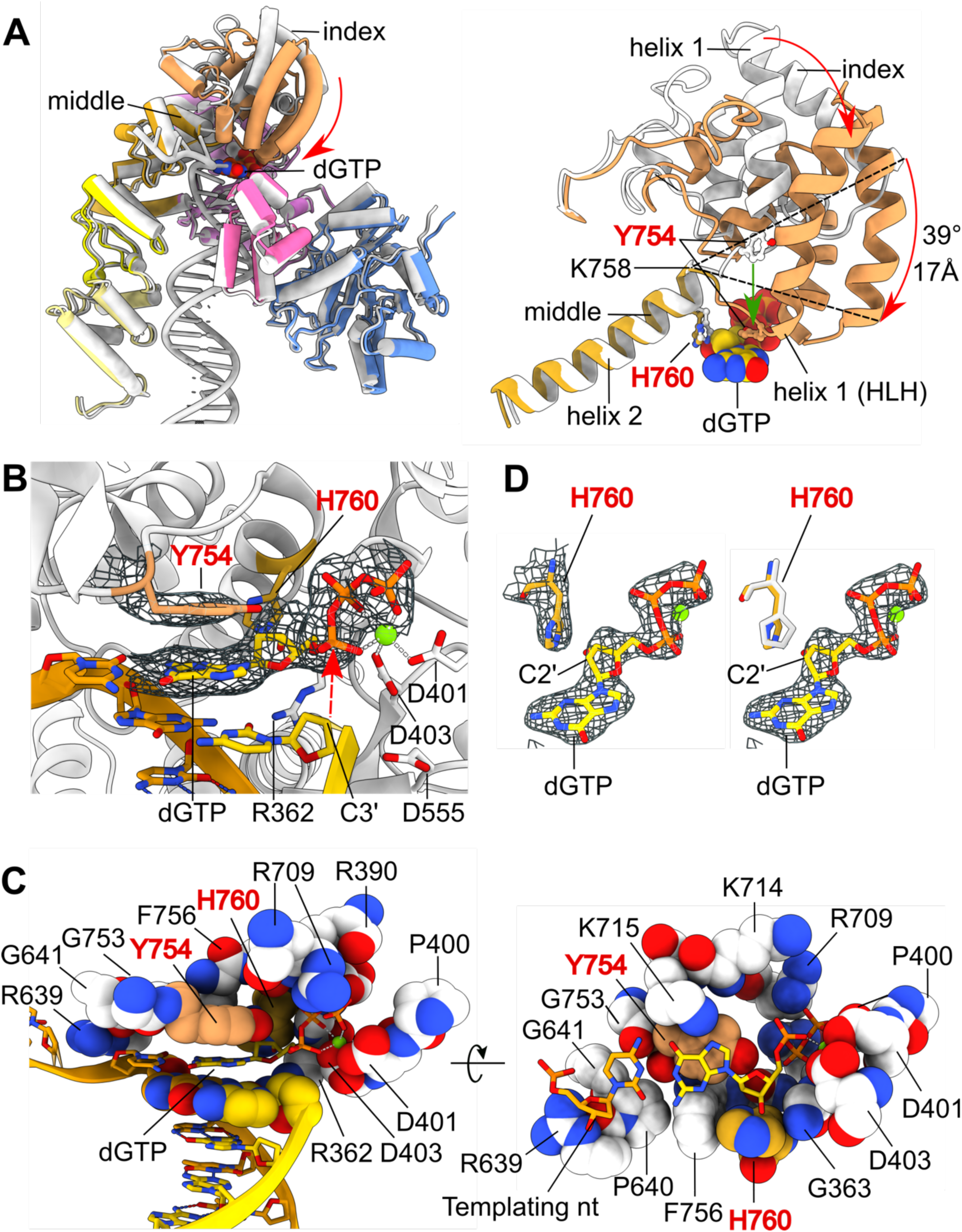
Polymerase active site and nascent base pair-binding pocket of α. (**A**), Superimposition of α^BC^ and α^TC^, showing conformational changes of α^BC^ induced by incoming nucleotide binding. Upon dGTP binding, the index finger undergoes a large 39° rotation pivoting around Lys758 to encapsulate dGTP, travelling up to 17 Å, while other domains of α are mostly unchanged. Domains of α^TC^ are coloured as Figure 3B. α^BC^ and DNA^TC^ are coloured white. Linker, OB and DNA^BC^ are omitted for clarity. Right, zoom-in view showing the index finger (sandy brown) rotates down onto dGTP (spheres) pivoting around Lys758 of the middle finger (goldenrod). This enables Tyr754 to close onto dGTP (green arrow). (**B**), Polymerase active site of α. The 3′-carbon of the terminal 2′,3′-dideoxycytidine monophosphate (ddCMP) on the primer is close to the α-phosphate of dGTP (red arrow). The phenolic ring of Tyr754 stacks on the base of dGTP and imidazole ring of His760 is close to the deoxyribose of dGTP. Densities of dGTP and Tyr754 are shown as blue meshes. (**C**), Nascent base pair-binding pocket of α in side view (left) and bottom view (right). Amino acid residues within 4 Å from the templating and incoming nucleotides are shown as spheres. DNA is shown as ribbon and sticks, except −1 nucleotides that form the bottom of the pocket are shown as spheres. Tyr754 (sandy brown) forms part of the roof of the binding pocket and stacks on dGTP, while His760 (goldenrod) forms part of the rear wall and blocks ribonucleotide binding. Lys714 and Lys715 that form the front wall are removed from the side view to allow viewing of the nascent base pair. (**D**), Zoom-in views of His760 and dGTP. Left, His760 of α^TC^ is close to the C2′ carbon of the deoxyribose moiety of dGTP and could clash with 2′-hydroxyl group if a rNTP was present. Right, the conformer of His760 in α^BC^ (white sticks) is incompatible with dNTP binding.

A dGTP sits in the nucleotide insertion site of α^TC^, base-pairing with the templating base (Figure 4B). The 3′-carbon of the terminal ddCMP on the primer is 4.6 Å from the α-phosphate of dGTP. If a dCMP instead of ddCMP is present, its 3′-hydroxyl group could be well poised for nucleophilic attack on the α-phosphate group of dGTP. A Mg^2+^ is in the Metal B site, coordinated by the β- and γ-phosphates of dGTP and the carboxylates of Asp401 and Asp403, while Mg^2+^ is missing from Metal A site, likely due to the lack of 3′-hydroxyl at the 3′-end of primer.

The roof and wall of the nascent base pair-binding pocket is formed by residues from the palm, index and middles fingers. Residues within 4 Å from dGTP are mainly located in three regions: the GS motif (Arg362, Gly363, Ser364) on Pβ1; Pβ2 that harbours catalytic Asp401 and Asp403; and a helix−loop−helix (HLH) motif comprised of the last helix of the index finger and the first helix of the middle finger (Tyr754, Phe756, Asn757, His760) (Figure 4C). Positively charged residues, Arg390 on the palm, and Arg709, Lys714 and Lys715 on the index finger are also within 4 Å from dGTP. Consistent with the structures, R390A mutation resulted in 27-fold lower catalytic efficiency partly through a significant increase in *K*_M_^26^. Additionally, residues Arg639, Pro640, Gly641 and Gly753 are within 4 Å of the templating nucleotide. Arg396, Lys553, and Arg710^25^, though slightly more distant (> 4 Å), can potentially interact with dGTP. For example, Lys553 forms a salt bridge with the last phosphate of the primer and a K553A mutation led to complete loss of polymerase activity^26^.

The GS motif is conserved in the Polβ-like β-NT superfamily^27^ and is implicated in nucleotide binding^1^. The motif is in close contact with dGTP. The backbone of Gly363 contacts the deoxyribose and β-phosphate of dGTP. Ser364 is close to both β- and γ-phosphates and forms two hydrogen bonds with dGTP. The side chain of Arg362 that originally points away from the active site in α^apo^ flips in and hydrogen bonds with O3′ of the deoxyribose moiety of dGTP. Arg362 is conserved in both α and PolC and R362A mutation resulted in complete loss of activity^26^.

The loop of the HLH motif is conserved in α and PolC and in a location similar to the nucleotide binding O-helix of Pol I polymerase that is implicated in replication fidelity and translocation^5^. In α^BC^, the HLH is distant from the active site. Upon binding of dGTP, the index finger rotates to close on dGTP. The C-terminal end of helix 1 becomes partially unwound and extended, allowing the conserved residues on the loop, Tyr754 and Phe756^5^, to form the roof above dGTP, and the phenyl group of Tyr754 to stack on the base of dGTP (Figure 4B,C, Movie 2). In *G. kaustophilus* PolC (3F2B), Tyr1269, an equivalent to Tyr754, also stacks on the base of the incoming nucleotide^18^. The stacking and the narrow binding pocket thus formed imposes steric restraints that strongly favours binding of a Watson-Crick base pair. Attempts to probe the role of Tyr754 through mutational analyses were limited by the inability to obtain viable *E. coli* colonies expressing Y754A or Y754F, indicating these α variants are highly toxic to *E. coli* cells.

Another conserved residue His760 on helix 2 of the HLH is also very close to dGTP, forming part of the rear wall of the pocket (Figure 4D). In α^apo^, His760 could sterically clash with a nucleotide in the active site. In α^TC^, His760 switches to another conformer that allows a dNTP to bind (Figure 4D, Movie 2). Its imidazole ring is close to the deoxyribose of dGTP and could clash with the 2′-hydroxyl group if an rNTP is present. In PolC, His1275, counterpart of His760, is similarly positioned^18^. Therefore, His760 can function as a steric gate to prevent incorporation of rNTPs^28^.

### The template overhang is expelled out of the central channel of α by incoming nucleotide

Next, we turned to computational simulations to study how the template overhang (extended to 10 nt for simulation, see Supplementary Methods) interacts with α. First, molecular dynamics simulation of PolIII^BC^ showed that the template overhang did not swing out of the central channel of α spontaneously due to electrostatic attraction to the positively charged residues on the surface of the channel (Supplementary Movie 3). Second, placing a dGTP at the active site of PolIII^BC^ quickly expelled the overhang out of the central channel (Supplementary Movie 4). However, within the 1-μs simulation, the overhang had not fully extended and yet established stable interactions with OB, still blocking the index finger from rotating downwards to close onto dGTP (Figure 5A). Third, simulation of PolIII^TC^ showed while the first 4 nt ahead of the templating nucleotide maintained contacts with the middle finger and OB, the remaining 5 nt did not bind stably to any parts of α, wiggling back and forth to interact with the index finger or OB, or protruding away from OB into space (Figure 5B−D, Movie 3). Contacts between this flexible part of the overhang and α continued to form and break. The interactions remained transient even when the entire overhang spanned over the surface of OB (Figure 5D). This is consistent with the 4 nt overhang seen in the cryo-EM maps and SPR results showing fast kinetics, whereby residues beyond +6 nt contribute much less to binding (Supplementary Figure 3 and Table 1). Taken together, our results suggest that interactions with the first 4−6 residues on an overhang ahead of the templating nucleotide may be critical for stabilising polymerase onto DNA for efficient polymerization (for detailed analysis see Supplementary Data 1 and 2). Residues beyond this point would likely associate with single-stranded DNA-binding proteins in the cell.

**Figure 5.**
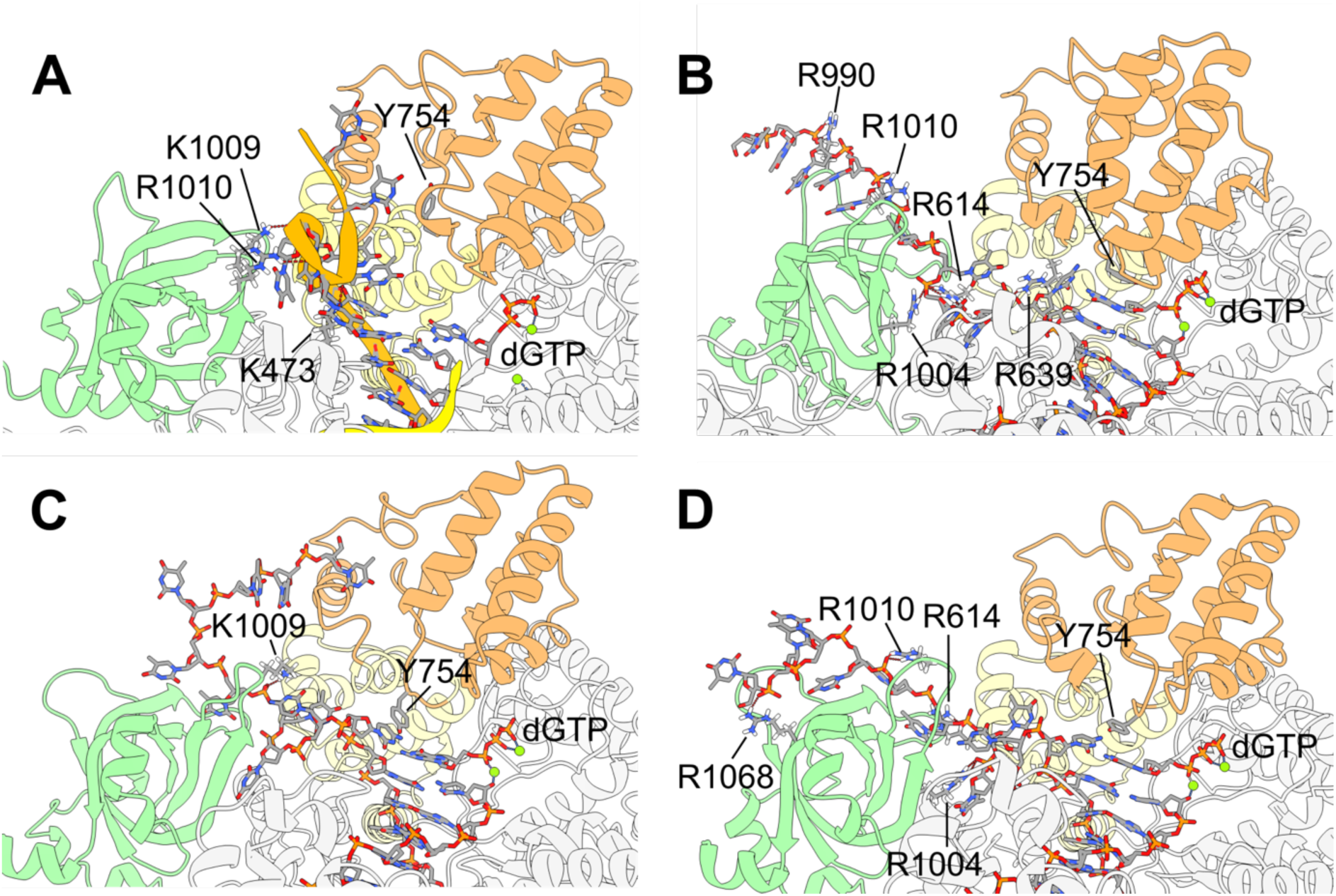
Representative frames of molecular dynamics simulations. (**A**), A representative frame of the molecular dynamics simulation of PolIII^BC^ with a dGTP manually placed in the active site of α (Supplementary movie 5). The ss template overhang displaced from the central channel of α has not yet fully extended (curled orange ribbon) in the 1-μs simulation, which interacts transiently with OB. The index finger of α is coloured sandy brown, middle finger pale goldenrod, OB light green, template strand orange and primer gold. Residues hydrogen bonding to the overhang are labelled and hydrogen bonds are indicated by red dash lines. (**B−D**), Representative frames of the molecular dynamics simulation of PolIII^TC^ (Movie 3), showing that the ss overhang beyond +4 nt continually shifts and interacts with different parts of α.

### Structural basis of proofreading by PolIII

To understand how PolIII HE removes mis-incorporated nucleotides, we determined the structures of αεθ·τ_C32_·β_2_ complexed with a p/t DNA containing a mismatch at the penultimate position (−1 nt) of the primer strand. This mismatch was expected to destabilise the complex, shifting the equilibrium towards the exonuclease state. Five structures were obtained, two are PolIII^BC^-like complexes with the index finger open and terminal nucleotide of primer sitting in the polymerase active site (PolIII^BC^*^1,2^) (Figure 6A), which account for 32.9% of total particles; and three are proofreading complexes with the 3′-end of the partially unwound primer strand located in the “active site” of the catalytically-dead ε^EL^ (PolIII^PR1-3^) (Supplementary Table 3, Supplementary Fig. 11).

**Figure 6.**
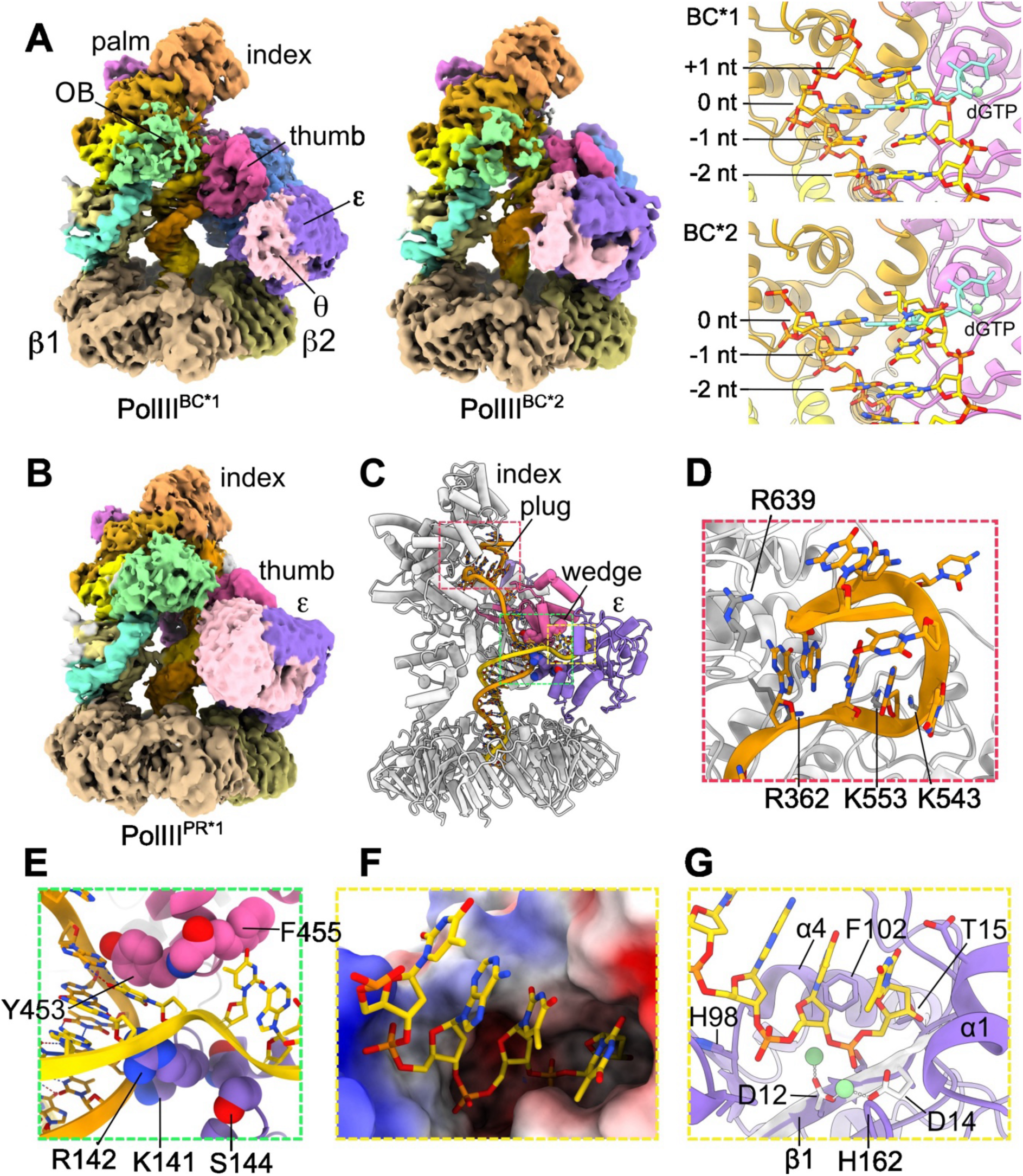
Structures of the *E. coli* PolIII^BC^-like and PolIII^PR^ complexes with a mismatched p/t DNA. (**A**), Cryo-EM maps of the *E. coli* PolIII^BC*1^ and PolIII^BC*2^ complexes. Right, zoom-in views of the active sites of PolIII^BC*1^ and PolIII^BC*2^. There is a 1 nt overhang (+1 nt) in PolIII^BC*1^ and no visible overhang in PolIII^BC*2^. dGTP in PolIII^TC^ (cyan) is also shown to indicate the location of nucleotide insertion site (0 nt) in α. Numbers in the right panel indicate positions of nucleotides on the template strand. (**B**), Cryo-EM map of PolIII^PR1^. (**C**), Structure of PolIII^PR1^. The thumb of α, ε, and the nascent and template strands are coloured hot pink, medium purple, gold and orange, respectively. The “wedge” on thumb and the tip of ε that stabilise the three-way junction are shown as spheres. (**D**), The template overhang forms a plug-like secondary structure in the central channel of α. Some residues that interact with the plug are labelled. (**E**), The three-way junction of DNA is stabilized by the “wedge” on thumb comprised of Tyr453, Gly454 and Phe455 and tip of ε (Loop Gly140−Ser144). Tyr453 may serve as a “separation pin”, interacting with both bases at junction and stacking on the base of the primer strand. Loop Gly140−Ser144 of ε interacts with the minor groove of dsDNA near junction. (**F**), The 3’-terminal nucleotide of the primer strand resides in the nuclease active site of ε. The surface of ε is coloured by electrostatic potentials. Blue indicates positively charged areas and red negatively charged. (**G**), Interactions of ε with the primer strand in the nuclease active site. ε is shown as cartoons and important residues involved in DNA interactions are shown as sticks. Crystal structure of the ε186·TMP complex (1J53) is superimposed with ε^PR^ to show the positions of the two catalytic metal ions (green spheres). Only the β1 strand of the ε186·TMP structure is shown as a white ribbon.

The structures of PolIII^BC^*^1,2^ are similar to PolIII^BC^. However, in contrast to six unpaired nucleotides (0 to +5 nt) seen in PolIII^BC^ (Figure 1C), there is only a 1-nt overhang in PolIII^BC^*^1^, and no visible overhang in PolIII^BC^*^2^ (Figure 6A, Supplementary Figs. 12 and 13). Moreover, the 3′-nucleotides of the primers reside in the polymerase active site of α^BC*^ (shifted by +1 nt in registry compared to PolIII^BC^) and do not properly pair with the templating nucleotide. Therefore, PolIII^BC^*^1,2^ resemble post-(mis)insertion but pre-proofreading intermediates containing a frayed DNA^25^. Compared to PolIII^BC^*^1^, DNA density in PolIII^BC*2^ is much weaker and α^BC^*^2^ looks more relaxed, suggesting weaker α−DNA interactions, which may be compensated by θ−OB contact.

The PolIII^PR1-3^ structures are similar to the 6.7 Å proofreading structure (*Eco*PolIII^PR^, 5M1S)^16^ with key differences on DNAs (Figure 6B,C). Comparing to PolIII^BC^*, DNA in PolIII^PR^ undergoes complex conformational changes, including rotation, retraction, tilting and partial unwinding, with the 3′-end of the primer strand moving ∼57 Å to the exonuclease active site (Movie 4). The template overhang lies in the central channel of α, coiling up to form a plug-like structure with several nucleotides stacking on one another (Figure 6D). The plug makes contacts with many residues on the index finger, such as, K543, R639, Pro640, Gly753, Phe756, and His760, and hydrogen bonds to Arg362 (Supplementary Fig. 14). The three-way junction is stabilized by a “wedge” on the thumb comprising Tyr453, Gly454 and Phe455, and the tip of ε (Figure 6E). Tyr453 poises like a “separation pin”, stacking on the base of the partially unwound primer strand at the junction. Phe455 stacks on the base of the first unpaired nucleotide and Gly454 hydrogen bonds to its phosphate group. Loop Gly140−Ser144 of ε interacts with the minor groove of ds DNA below the junction. Gly140 and Lys141 contact the backbone of the template strand and Arg142 hydrogen bonds to O4′ of the deoxyribose of the nucleotide of the nascent strand at the junction. The ds DNA passes through the β_2_ ring at ∼30° with minor contact to loop Leu21−Arg24 of the β1 subunit opposite to ε. The middle section of the ss template does not interact with α much, thus α^PR^ is more relaxed than α^BC^*.

There are four unpaired nucleotides on the primer strand, rather than three observed in *Eco*PolIII^PR^ (5M1S) (Figure 6F). The four unpaired bases line up and stack on one another. The terminal nucleotide positions in the nuclease active site like TMP_A_ in the ε186·TMP complex (1J53)^23^, while the penultimate nucleotide sits deeper than TMP_B_. The terminal nucleotide is flanked by the β1 strand of ε, the following loop and the α1 helix (Figure 6G). It interacts with Thr13, Thr15, Gly17, Val65, Phe102 and His162 (Supplementary Fig. 14). The penultimate nucleotide is close to α4 and interacts with His98, Phe102, Leu145 and His162. Catalytic residues Asp12(Ala) and Glu14(Ala) on β1, and His162 that was suggested to act as a general base to deprotonate the active site nucleophile^23^ are close to the phosphate group of the terminal nucleotide. Due to the alanine replacements of Asp12 and Glu14 in ε^DL^, the two metal ions that coordinate TMPs in ε186·TMP structure are absent. If present, the two metal ions would be well positioned to contact the last phosphate group (Figure 6G). Positions of the third and fourth nucleotides are shallower, sitting at the entrance of the nuclease active site. The phosphate group of the third nucleotide is hydrogen bonded to Arg135 and Asn143.

## DISCUSSION

The high-resolution structures of the *E. coli* αεθ·β_2_·DNA binary, ternary and proofreading complexes establish the structural basis of nucleotide selection, incorporation, and proofreading by PolIII, revealing how PolIII proceeds through the catalytic cycle to achieve highly processive and high-fidelity DNA replication (Figure 7).

**Figure 7.**
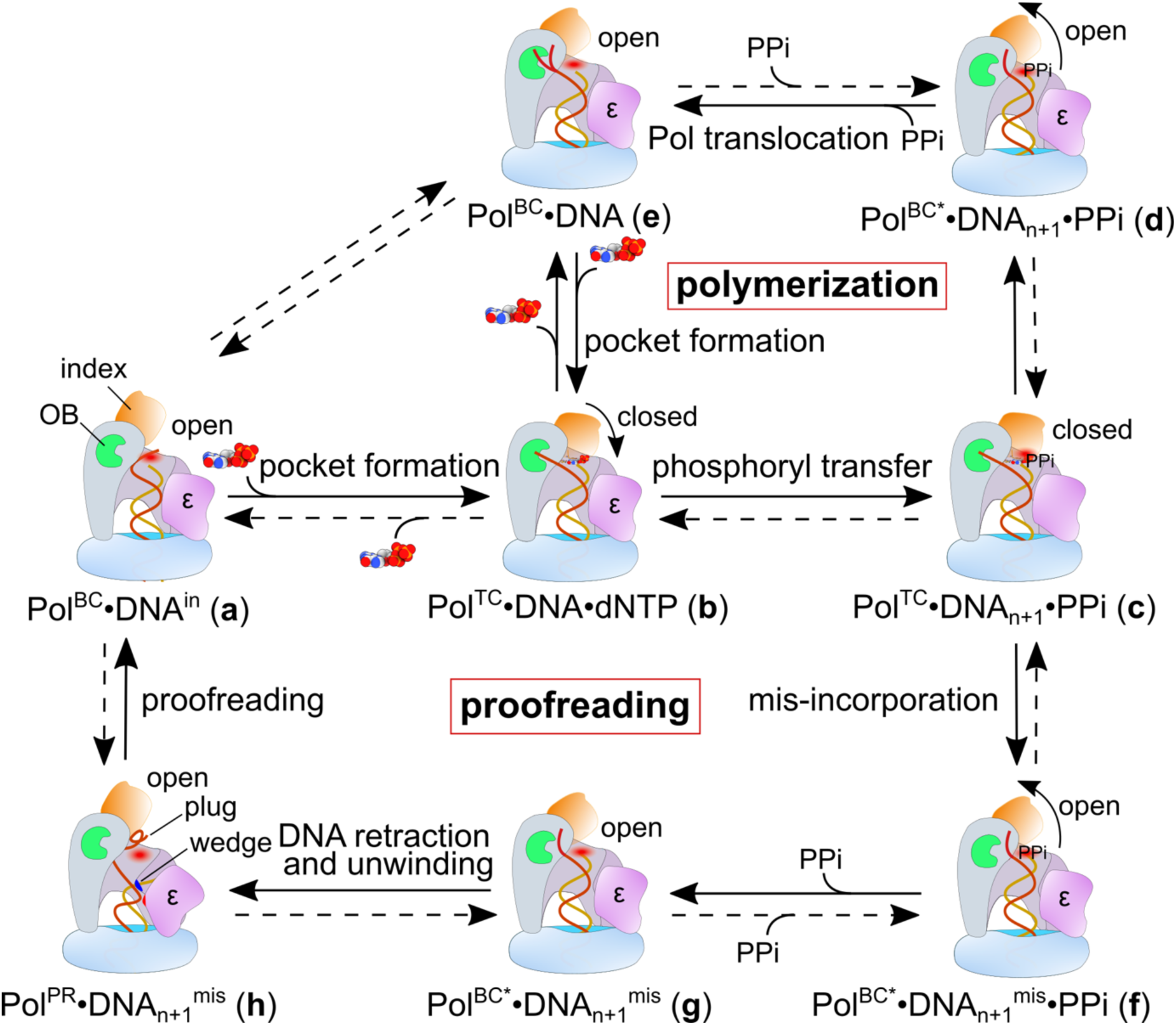
Pathways of DNA synthesis and proofreading by PolIII. First, α binds to β_2_-loaded primer-template DNA, undergoing significant conformational changes to adapt to DNA binding (**a**, PolIII^BC^·DNA^in^). However, the index finger of α remains “open”. The ss template overhang sits in the central channel of α, which does not swing out to the side path spontaneously. Binding of an incoming nucleotide in the polymerase active site (red halo) expels the overhang to the side path whereby it contacts OB, allowing the index finger to rotate towards and Tyr754 to stack onto incoming nucleotide, and the nascent base pair-binding pocket to form (**b**, PolIII^TC^·DNA·dNTP). Then, the phosphoryl transfer reaction proceeds, resulting in nucleotide incorporation and production of a pyrophosphate (**c**, PolIII^TC^·DNA_n+1_·PPi). Subsequently, the index finger opens, the polymerase subunit may relax its hold on DNA, and the template overhang may dissociate from OB temporarily, which may resemble the PolIII^BC^-like complex we obtained with a mismatched DNA (**d**, PolIII^BC*^·DNA_n+1_·PPi). The pyrophosphate then dissociates, and the polymerase moves forward on DNA by 1 nt, emptying the nucleotide insertion site (**e**, PolIII^BC^·DNA). The template overhang may either contact OB or dissociate from it but remain outside the centre channel during processive polymerization (**e**). An incoming nucleotide can quickly bind and a new cycle begins (**b**). Non-complementary nucleotides failing to base pair will dissociate (**b** to **e**). In rare cases, a mismatched dNTP may pass through check points and a binding pocket form (**b**). However, the configurations of the binding pocket would likely deviate from normal and mis-align the dNTP with the 3′-OH of primer that will substantially slow down phosphoryl transfer reaction, resulting in high probability of nucleotide dissociation (**b** to **e**). On the other hand, binding of a complementary rNTP would be blocked by His760. If phosphoryl transfer does occur with a wrong nucleotide (**c**), the mismatch would result in DNA distortion and weakened polymerase–DNA interactions (**f** then **g**, PolIII^BC*^·DNA_n+1_^mis^). Subsequently, DNA retracts from the polymerase active site and unwinds, transferring the mismatch to the exonuclease active site of ε for proofreading; while the ss template strand forms a “plug” to anchor in the central channel of α (**h**, PolIII^PR^·DNA_n+1_^mis^). Once the mismatch is removed, the processed nascent strand returns to the polymerase active site (**a**). Unnecessary (excessive) cleavage of the nascent strand may be prevented by timely dissociation of DNA from ε due to weakened DNA−ε/α interactions upon cleavage of the terminal nucleotide. Dashed arrows indicate the reverse reactions may be possible but less likely.

### Mechanisms of nucleotide selection by PolIII

High-fidelity DNA synthesis by polymerases relies on: 1, Watson-Crick geometry of a nascent base-pair. All high-fidelity DNA polymerases possess a narrow base-pair-binding pocket, in which only a Watson-Crick base pair can properly bind^29–32^. 2, the inefficiency of polymerases to incorporate mismatched nucleotide due to mis-alignment of the 3′-hydroxyl group of primer with the α-phosphate of the incoming nucleotide^33^. 3, a 3′−5′ exonuclease to remove wrongly inserted nucleotides^31,34^.

DNA polymerases typically require rotation of fingers to close on an incoming nucleotide and form a narrow binding pocket for nucleotide selection^35,36^, *e. g.*, the Pol I DNA polymerases use a bridge helix (O-helix) to close onto the nascent base-pair^37,38^. Likewise, the index finger of α undergoes a large rotation to close onto the nascent base-pair, forming the roof and main wall of the nascent base-pair-binding pocket (Figure 4, Movie 2). During the rotation, the C-terminal end of helix 1 of the HLH motif becomes partially unwound and extended, allowing the conserved Tyr754 to stack its phenyl ring on top of the nucleoside base of the incoming dNTP. The narrow binding-pocket which forms can only optimally fit a Watson-Crick base pair, positioning the incoming nucleotide for nucleophilic attack by the 3′-hydroxyl of the primer. In rare cases, mismatched dNTPs may pass through check points and a binding pocket still forms. However, such a binding pocket would likely deviate from normal configurations and mis-align the incoming dNTP with the 3′-OH of primer that will substantially slow down phosphoryl transfer reaction, resulting in much higher probability of nucleotide dissociation over integration (Figure 7).

In *E. coli* cells, the concentrations of rNTPs can be 100-fold higher than the corresponding dNTPs. Thus, rNTPs are often incorporated into DNA with frequencies three orders of magnitude higher than the wrong dNTPs^39,40^. Our structures with incoming nucleotide reveal that the imidazole ring of His760 is in close proximity to the 2′ carbon of the incoming dNTP’s deoxyribose and would sterically clash with a hydroxyl group at this position (Figure 4D), explaining how rNTPs are selected against. The mechanism appears universal. For example, Glu710 of *E. coli* Pol I (Family A) and Tyr254 of ϕ29 Pol (Family B) also act as steric gates to the 2′-hydroxyl of rNTPs^31,41,42^. Like α, *G. kaustophilus* PolC (3F2B) also possesses a similar HLH with Tyr1269, an equivalent of αTyr745, stacking on the incoming nucleotide; and His1275 similarly positioned as His760. Therefore, the structural basis of nucleotide selection is conserved in these two distinct branches of Family C DNA polymerases, although Gram-negative and positive bacteria diverged more than a billion years ago^43^.

### Proofreading by PolIII and potential mechanisms preventing unnecessary trimming of DNA

Rapid and efficient removal of mis-incorporated nucleotides by the 3′−5′ exonuclease ε provides another mechanism to ensure highly processive high-fidelity DNA replication by PolIII. Mechanisms preventing unnecessary trimming of DNA and ensuring prompt resumption of DNA synthesis are also important. Our structures captured several high-resolution snapshots of the proofreading intermediates. The PolIII^BC*1,2^ structures show that a mismatch resulted in distortion and fraying of DNA and opening of the index finger (Figure 6A). Subsequently, DNA retracts from the polymerase active site and unwinds until the 3′-end of the partially unwound primer strand is transferred to the nuclease active site of ε, while the ss template strand forms a plug in the central channel of α (PolIII^PR^) (Figure 6D, Movie 4). The three-way DNA junction is stabilised by a “wedge” on thumb and a tip of ε and the 3′-end of the primer strand is secured in the nuclease active site, positioning the last phosphodiester bond for cleavage by the two metal ions in the active site (Figure 6G). Once the mismatch is removed, DNA−α/ε interactions are likely weakened due to the loss of the terminal nucleotide, allowing the processed primer strand to dissociate from ε and return to the polymerase active site to re-base-pair with the template strand. Alternatively, ε may continue to remove more unwound nucleotides. However, this would require further unwinding of the ds DNA to optimally position the new terminal nucleotide for cleavage. Therefore, cleavage of the terminal nucleotide (or more) itself can serve as a built-in mechanism that prevents excessive cleavage and ensures quick resumption of DNA synthesis.

### Roles of OB

OB of PolIIIα was proposed to bind and guide the template strand into the polymerase active site^17^. In PolIII, the Linker connecting the little finger and OB is rigid so that it restrains OB close to the side path, consistent with the proposed role of OB. At least a 4-nt overhang is needed to reach OB. This is consistent with our SPR results that incoming nucleotide did not increase the binding affinity of α to DNA with a 2-nt overhang (*i.e.*, *K*_D1_/*K*_D_^app^ ≅ 1), but significantly increased binding affinity to DNA with a 4-nt overhang or longer (Table 1). The contributions become less significant for overhangs longer than 6 nt, for which *K*_D1_/ *K*_D_^app^ = 9.3. In our structures, no density of ssDNA beyond 4 nt is seen, suggesting that template overhang is not tightly bound to OB. Molecular dynamics simulations show while the first 4 nt on the overhang maintains contacts with OB, the remaining nucleotides shifted continually, either interacting with the middle finger or OB; or protruding away from OB (Figure 5B−D, Movie 3). The transient nature of DNA−OB interactions, rather than tight static binding, is well-suited for high-speed DNA synthesis. Different from PolIIIα, OB in PolC (3F2B)^18^ is located opposite to the side path (Supplementary Fig. 1E). It was suggested that OB^PolC^ may interact with ss template ∼15−20 nt ahead of the polymerase active site. Although OB in such a position cannot directly guide the template strand to active site, it may stabilize and prevent it from swinging back into the central channel of PolC during processive synthesis.

In conclusion, this work defines the molecular mechanism by which bacterial DNA polymerase III selects, incorporates, and proofreads nucleotides to achieve rapid, processive, and highly faithful genome replication. The combination of high-resolution structural snapshots, biochemical analyses, and molecular dynamics simulations captures the complete nucleotide incorporation cycle at near-atomic resolution (Figure 7), providing a long-sought mechanistic framework for bacterial DNA replication and a foundation for understanding the conserved principles that govern replication fidelity across cellular life.

## METHODS

### Plasmids and strains

*E. coli* strain AN1459 was used for plasmid construction and strains BL21(λDE3)*recA^−^*, BL21 (λDE3)/pLysS and BL21-AI for protein expression. Plasmid pZX2218 encoding the α^E612K/QLDLF^ (α^EL^) variant was constructed by replacing the XhoI−RsrII restriction fragment of pKO1538 (encoding α^E612K^) with the corresponding fragment of pSJ1392 encoding α^QLDLF^. Plasmid pNL2232 encoding the ε^D12A/E14A/QLSLPL^ (ε^DL^) variant was constructed by replacing the AflII−HindIII fragment of pKO1429 (encoding ε^D12A/E14A^) with that of pSJ1446 (encoding ε^QLSLPL^)^11^. Plasmids encoding α^Y754A^ or α^Y754F^ in pET-29b(+) were obtained from Twist Bioscience.

### Overproduction and purification of proteins

BL21-AI cells harbouring pZX2218 and pNL2232 were used for overproduction of α^EL^ and ε^DL^, respectively. Briefly, 7 ml of overnight culture in Luria-Bertani medium (LB) was inoculated into 1 l of LB containing 100 mg l^−1^ ampicillin (totally 12 l for α^EL^, 4 l for ε^DL^). The bacteria were grown at 37°C until OD_600_ ∼0.6. Protein overproduction was induced by adding L-(+)-arabinose to 0.2% (w/v). The bacteria were then grown at 30°C (α) or 37°C (ε) for 4 h. Cells were harvested by centrifugation and cell pellets stored at −80°C. α^E612K^ and α^EL^ were purified as per Lewis^44^, ε^DL^ and the PolIII α^EL^ε^DL^θ core as per Jergic^11^, τ_3_δδ’ψχ per Tanner^45^ and τ_C32_ per Monachino^22^.

### Polymerase–DNA interaction analysis by surface plasmon resonance (SPR)

Interactions between α^E612K^ and wild-type α with p/t DNAs were measured at 25°C in SPR buffer: 30 mM Tris, pH 7.6, 50 mM NaCl, 12 mM MgCl_2_, 0.5 mM EDTA, 0.5 mM DTT containing 0.005% surfactant P20. First, biotinylated primer 5′-bio-AAAACGAAAAATAAGTATGTTGTAACTAAAG(ddC) (32-mer) was immobilized onto the streptavidin coated (SA) chip surface. Then, a template strand from the set of oligos 5′-(C)_x=3−15_GCTTTAGTTACAACATACTTATTTTTCGTTTT (500 nM) was injected to hybridize with primer. Interactions of α^E612K^ and wild-type α with p/t DNA were mostly measured by sequential injections of serially diluted samples in SPR buffer for 30 s at 20 and 100 μl/min, respectively, followed by dissociation over 60 s.

The sensorgrams were subtracted from unmodified ligand flow path using BIAcore software and zero subtracted using MATLAB. Equilibrium binding parameters (dissociation constant, *K*_D_, and apparent dissociation constant, *K*_D_^app^) were determined using a 1:1 steady state affinity (SSA) binding model:

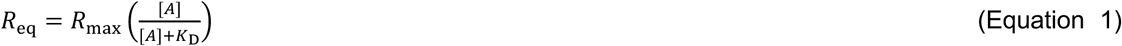

where *R*_max_ corresponds to the response when all the immobilized ligands (p/t DNA) on the surface are saturated with the analyte A (polymerase), *K*_D_ is the dissociation constant, and [A] is the concentration of analyte. *R*_eq_ represents equilibrium binding response upon interaction with surface immobilized DNA. A thorough description of SPR analysis is included in Supplementary Information.

### Sample preparation for single-particle cryo-EM

DNA for the αεθ·τ_C32_·β_2_·DNA binary complex was prepared by annealing Oligo 1 (5′-GAGATAGTTACAACATACGATCG) (Integrated DNA Technologies, IDT) to Oligo 2 (5′-T_10_-CGATCGTATGTTGTAACTATCTC). DNA for the αεθ·τ_C32_·β_2_·DNA·dGTP ternary complex was prepared by annealing Oligo 3 [5′-CCAAATAAGTATGTTGTAACTAAAG(ddC)] (TriLink Biotechnologies, PAGE purified) to Oligo 4 (5′-T_48_-CCGCTTTAGTTACAACATACTTATTTGG) (IDT). DNA for the αεθ·τ_C32_·β_2_·DNA^mismatch^ binary-like pre-proofreading and proofreading complexes was prepared by annealing Oligo 5 (5′-CTTAGTTACAACATACTTATT) to Oligo 6 (5′-CAGACATCCGTAGACTTCGGTCATTAAGTAT GTTGTAACTAAG).

To prepare the αεθ·τ_C32_·β_2_·DNA sample, 20 μl of 52.3 μM α^EL^ε^DL^θ was mixed with 10.5 μl of 150 μM p/t DNA, 10.4 μl of 130 μM β_2_, 16.2 μl of 84 μM τ_C32_, 2 μL of 200 mM MgCl_2_, and dialysed twice at 4°C against 250 ml of 30 mM Na.HEPES pH 7.5, 5 mM MgCl_2_, 2 mM dithiothreitol, 0.25 mM EDTA, 2% glycerol. For the ternary complex, dGTP was added to the dialysate to a final concentration of 1 mM. Samples (3 μl) were applied onto Quantifoil UltrAuFoil grids (R1.2/1.3) that were blotted for 4.0 s at 8°C and zero force, then plunged into liquid ethane using a FEI Vitrobot Mark IV.

### Electron microscopy data acquisition, image processing and model building

Micrographs were collected using a Thermo Fisher (FEI) Titan Krios G3i microscope at 300 kV equipped with a Gatan K3 camera in electron counting mode. Micrographs were acquired at a nominal defocus range of −0.4 to −1.2 mm with a magnification of 0.83 Å/pixel, a total dose of 65 e^−^/Å^2^ accumulated over 6.7 s and fractionated across 100 frames. A subset of micrographs was collected at a 30° tilted angle to minimize preferred orientation.

Data were processed in CryoSPARC v4.5.3^46^. Frames were corrected for drifting and aligned. CTF parameters were estimated using dose-weighted images and micrographs with estimated resolutions lower than 4.5 Å were excluded. Particles were first picked with blob-picker. The particles were subjected to 2D classification and selected 2D classes were used for Template Picking. Extracted particles were subjected to rounds of 2D classification and only averages showing high-resolution features were retained. Selected particles were subjected to two rounds of 3D classification by Ab- Initio Reconstruction followed by Heterogenous Refinement. Particles of the best polymerase classes were selected and further classified into distinct 3D classes by 3D Classification. Particles of each individual class were subjected to Non-uniform Refinement, followed by Reference-based Correction and a final Non-uniform Refinement.

For model building, structures of the *E. coli* apo α (2HNH)^5^, εθ (2XY8)^23^ and β_2_ (1MMI)^47^ were placed into the density map using Phenix.dock_in_map^48^. DNA was built into the map in Coot^49^. The resulting structure was refined in Phenix.real_space_refine. After inspection and model building in Coot, dGTP and MgCl_2_ were included in the model. The model was subjected to a cycle of rebuilding in Coot and REFMAC refinement, followed by a final cycle of refinement in Phenix.real_space_refine. Structure analyses used ChimeraX^50^ and the PDBePisa server^51^. Figures and movies were prepared using ChimeraX^50^.

## Acknowledgements

We thank Enrico Monachino, Nan Li, Kiyoshi Ozawa, Allen Lo, and Nick Horan for gifts of proteins or plasmids. This work was supported by the Australian Research Council (DP180100805 to A. J. O. and N. E. D.; DP210100365 to H. G., P. J. L., G. T., A. J. O. and N. E. D.; CE230100021 to H. Y.).

## Author contributions

Conceptualization, Methodology, Z.-Q. X., S. J., H. G., P. J. L., G. T., A. J. O., N. E. D.; Investigation: Z.-Q. X. (EM and structure analysis), S. J. (SPR, replication assay), Q. Z., H. B. Y. (Computational simulation), A. J. O. (model building), S. H. J. B., J. C. B. (EM support); Resources, Z.-Q. X., S. J., J. L.; Original Draft and Visualization, Z.-Q. X.; All authors have read the manuscript.

## Additional information

## Data availability

Cryo-EM maps and atomic coordinates to be deposited at wwPDB.

**Movie 1. Conformational changes of α upon DNA binding.** Morphing of structures from apo α to α^BC^, showing how α undergoes conformational changes on DNA binding. PHP, palm, thumb, and the index, middle, ring and little fingers of α are coloured cornflower blue, orchid, hot pink, sandy brown, goldenrod, yellow, and khaki, respectively.

**Movie 2. Conformational changes of the helix-loop-helix motif of** α **on dGTP binding.** Morphing of structures from α^BC^ to α^TC^, showing how α, especially the HLH motif on the index finger, undergoes significant conformational changes on dGTP binding. Tyr754 and His760 are shown as balls and sticks and dGTP is shown as spheres.

**Movie 3. The ss template overhang does not bind tightly to OB.** The movie shows the trajectory of the 1-μs molecular dynamics simulation of PolIII^TC^. During the simulation, while the first 5 nt of the ss overhang (in rainbow colours) downstream the ss-ds DNA junction (0−+4 nt) maintains contacts with OB and fingers, the remaining nucleotides (+5−+9 nt) does not bind tightly to any parts of α. They constantly shift, wiggling back to interact with the middle finger or OB, or protruding away from OB.

**Movie 4. Conformational changes during transition from PolIII^BC*^ to PolIII^PR^**. Morphing of structures from PolIII^BC*1^ to PolIII^BC*2^, then PolIII^PR^, showing conformational changes upon insertion of a mismatched nucleotide.

