## Supplementary Information for "Structural basis of the nucleotide incorporation cycle of bacterial DNA polymerase III"

### Structural basis of high-processivity high-fidelity DNA replication by *Escherichia coli* DNA polymerase III

**This document includes:**

Supplementary Tables 1–3

Supplementary Figures 1–14 and figure captions

Supplementary Movies captions

Supplementary Methods

Supplementary Data 1 – Model for fitting SPR data obtained in the presence of complementary nucleotide dGTP

Supplementary Data 2 – Summary and extended discussions of SPR studies on $\alpha$ binding to p/t-DNAs

Supplementary References

**Supplementary Table 1 | Details of cryo-EM data collection, refinement and validation of the *E. coli* PolIII αεθ•β₂•DNA binary complexes**

| | $PolIII^{BC1}$ | $PolIII^{BC2}$ | $PolIII^{BC3}$ |
|---|---|---|---|
| **Data collection and processing** | | | |
| microscope | Titan Krios G3 | | |
| voltage (kV) | 300 | | |
| electron dose ($e^{-}/Å^2$) | 65 | | |
| dose rate ($e^{-}/Å^2$/fraction) | 1 | | |
| detector | K3 | | |
| defocus range (μm) | 0.7–1.2 | | |
| calibrated pixel size (Å) | 0.83 | | |
| micrographs (No.) | 5949 (normal) and 1730 (tilted at 30°) | | |
| total extracted particles | 1811k | | |
| final particles | 338.0k | 335.1k | 310.7k |
| symmetry imposed | C1 | C1 | C1 |
| resolution (Å) | 2.54 | 2.47 | 2.47 |
| **Model-to-Map Fit** | | | |
| CC_mask | 0.77 | 0.78 | 0.79 |
| CC_volume | 0.77 | 0.78 | 0.78 |
| **Refinement** | | | |
| RMS deviations | | | |
| – bond lengths (Å) | 0.003 | 0.003 | 0.003 |
| – bond angles (°) | 0.585 | 0.626 | 0.633 |
| Ramachandran (%) | | | |
| – outliers | 0.05 | 0.31 | 0.21 |
| – allowed | 5.38 | 4.48 | 5.47 |
| – favoured | 94.57 | 95.21 | 94.32 |
| rotamer outliers (%) | 2.36 | 2.42 | 2.48 |
| Molprobity score | 2.15 | 2.05 | 2.14 |
| Clashscore | 9.28 | 7.77 | 8.35 |
| Q-score | 0.425 | 0.475 | 0.459 |

**Supplementary Table 2 | Details of cryo-EM data collection, refinement and validation of the *E. coli* PolIII αεθ•β$_2$•DNA•dGTP ternary complexes**

| | PolIII$^{TC1}$ | PolIII$^{TC2}$ | PolIII$^{TC3}$ | PolIII$^{TC4}$ |
|---|---|---|---|---|
| **Data collection and processing** | | | | |
| microscope | Titan Krios G3 | | | |
| voltage (kV) | 300 | | | |
| electron dose ($e^-/Å^2$) | 65 | | | |
| dose rate ($e^-/Å^2$/fraction) | 1 | | | |
| detector | K3 | | | |
| defocus range (μm) | 0.7–1.2 | | | |
| calibrated pixel size (Å) | 0.83 | | | |
| micrographs (No.) | 5718 (normal) and 4272 (tilted at 30°) | | | |
| total extracted particles | 2043k | | | |
| final particles | 351.3k | 369.3k | 421.6k | 379.8k |
| symmetry imposed | C1 | C1 | C1 | C1 |
| resolution (Å) | 2.61 | 2.62 | 2.51 | 2.52 |
| **Model-to-Map Fit** | | | | |
| CC_mask | 0.85 | 0.84 | 0.83 | 0.84 |
| CC_volume | 0.84 | 0.83 | 0.83 | 0.83 |
| **Refinement** | | | | |
| RMS deviations | | | | |
| – bond lengths (Å) | 0.003 | 0.005 | 0.003 | 0.003 |
| – bond angles (°) | 0.616 | 0.643 | 0.580 | 0.579 |
| Ramachandran (%) | | | | |
| – outliers | 0.09 | 0.14 | 0.14 | 0.15 |
| – allowed | 5.78 | 4.48 | 4.51 | 4.08 |
| – favoured | 94.12 | 95.38 | 95.35 | 95.77 |
| rotamer outliers (%) | 2.47 | 1.90 | 0.90 | 1.69 |
| Molprobity score | 2.09 | 1.96 | 1.62 | 1.80 |
| Clashscore | 7.21 | 7.72 | 5.49 | 6.04 |
| Q-score | 0.441 | 0.451 | 0.507 | 0.502 |

**Supplementary Table 3 | Details of cryo-EM data collection, refinement and validation of the *E. coli* PolIII αεθ•β₂•DNA proofreading complexes**

| | PolIII$^{BC*1}$ | PolIII$^{BC*2}$ | PolIII$^{PR1}$ | PolIII$^{PR2}$ | PolIII$^{PR3}$ |
|---|---|---|---|---|---|
| **Data collection and processing** | | | | | |
| microscope | Titan Krios G3 | | | | |
| voltage (kV) | 300 | | | | |
| electron dose ($e^{-}$/Å$^2$) | 65 | | | | |
| dose rate ($e^{-}$/Å$^2$/fraction) | 1 | | | | |
| detector | K3 | | | | |
| defocus range (μm) | 0.7–1.2 | | | | |
| calibrated pixel size (Å) | 0.83 | | | | |
| micrographs (No.) | 6078 (normal) and 4583 (tilted at 30°) | | | | |
| total extracted particles | 5096k | | | | |
| final particles | 431.0k | 540.5k | 344.7k | 400.4k | 329.2k |
| symmetry imposed | C1 | C1 | C1 | C1 | C1 |
| resolution (Å) | 2.42 | 2.73 | 2.48 | 2.58 | 2.37 |
| **Model-to-Map Fit** | | | | | |
| CC_mask | 0.84 | 0.83 | 0.84 | 0.79 | 0.87 |
| CC_volume | 0.83 | 0.82 | 0.84 | 0.79 | 0.86 |
| **Refinement** | | | | | |
| RMS deviations | | | | | |
| – bond lengths (Å) | 0.003 | 0.003 | 0.004 | 0.004 | 0.003 |
| – bond angles (°) | 0.574 | 0.551 | 0.530 | 0.592 | 0.526 |
| Ramachandran (%) | | | | | |
| – outliers | 0.05 | 0.05 | 0.10 | 0.10 | 0.26 |
| – allowed | 4.80 | 5.05 | 3.70 | 5.64 | 2.71 |
| – favoured | 95.47 | 94.89 | 96.19 | 94.25 | 97.03 |
| rotamer outliers (%) | 1.61 | 1.24 | 0.99 | 2.86 | 1.43 |
| Molprobity score | 1.86 | 1.67 | 1.43 | 2.15 | 1.43 |
| Clashscore | 7.10 | 4.81 | 3.83 | 7.48 | 3.54 |
| Q-score | 0.494 | 0.475 | 0.531 | 0.485 | 0.553 |

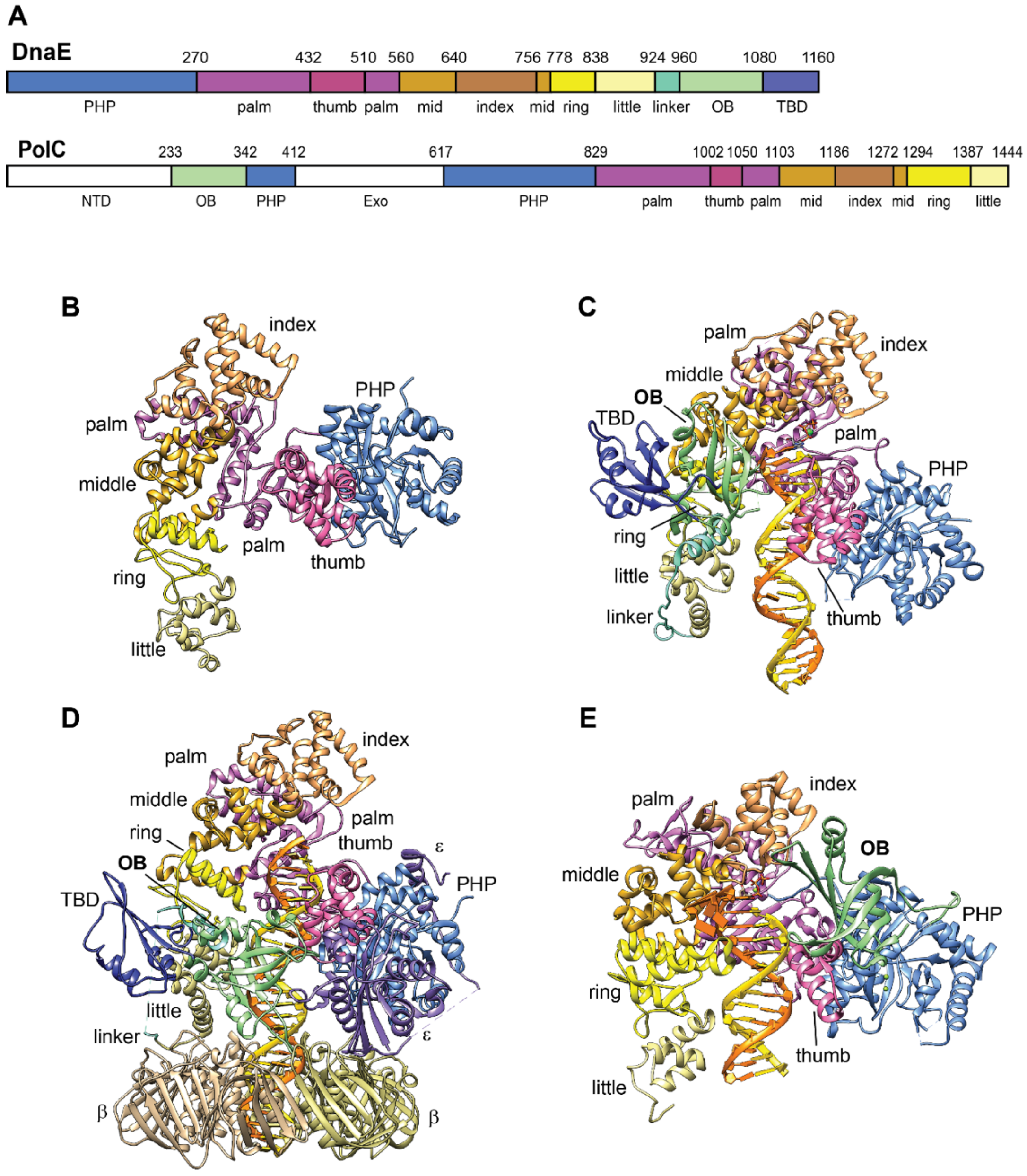

**Supplementary Figure 1**. **Structures of C-family DNA polymerases.** (**A**), Domain organization of DnaE (2HNH) and PolC (3F2B). DnaE (PolIII$\alpha$) is comprised of several domains (from N-terminus to C-terminus), including the PHP (polymerase and histidinol phosphatase) domain, palm, thumb, fingers (further divided into the index, middle, ring and little fingers), a linker bridging the little finger and OB (oligonucleotide/oligosaccharide binding) domain, OB and TBD ($\tau$-binding domain), which are coloured cornflower blue, orchid, hot pink, sandy brown, goldenrod, yellow, khaki, aquamarine, light green and light blue, respectively. Note, PolC possesses an extra N-terminal domain and a 3′ to 5′ exonuclease domain, which are coloured white. The domains are also arranged in a different order to DnaE. (**B**), Structure of the large N-terminal fragment of *E. coli* apo PolIII$\alpha$ (2HNH, 2.3 Å) lacking the Linker, OB and TBD. Individual domains are coloured as (**A**). (**C**), Structure of the *Thermus aquaticus* PolIII$\alpha$•DNA•dATP ternary complex (*Taq*PolIII$^{TC}$, 3E0D, 4.6 Å). The primer strand of DNA is coloured gold and template strand orange. (**D**), Structure of the *E. coli* $\alpha\varepsilon\theta$•$\beta_2$•$\tau_{C16}$•DNA binary complex (*Eco*PolIII$^{BC}$, 5FKV, 8 Å). $\varepsilon$ is coloured medium purple, and the $\beta$ subunits dark khaki and tan, respectively. $\tau_{C16}$ is omitted for clarity. Note the significant shift of *Eco*$\alpha$ OB towards little finger and clamp compared to the OB location in *Taq*$\alpha^{TC}$ (**C**). (**E**), Structure of the *Geobacillus kaustophilus* PolC•DNA•dGTP ternary complex (3F2B, 2.4 Å).

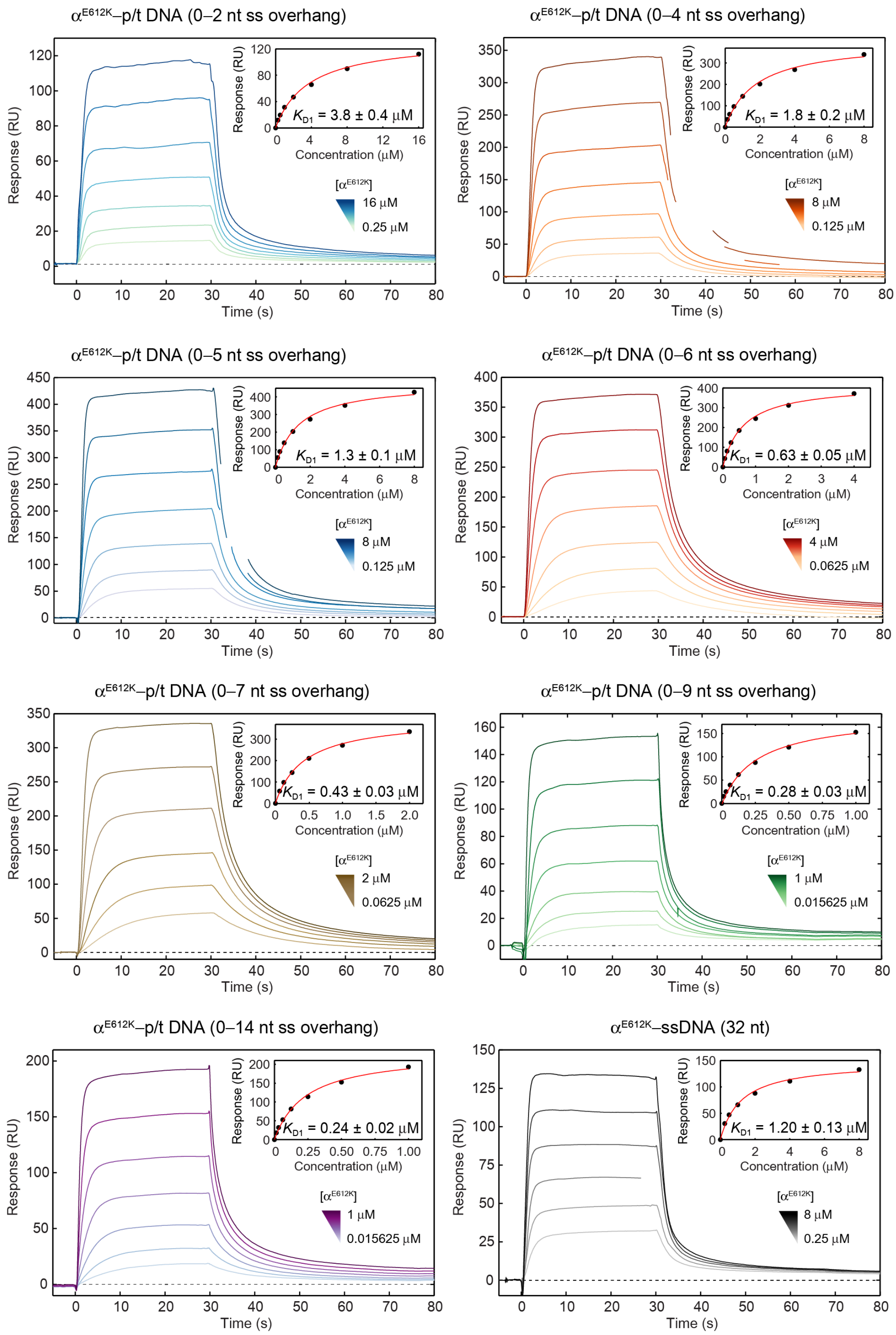

**Supplementary Figure 2. Binding of $\alpha^{E612K}$ to p/t DNA increases with increased number of residues on template overhang (for up to ~10 nt).** SPR sensorgrams showing association (30 s) and dissociation of $\alpha^{E612K}$ to and from a p/t DNA or ss DNA over a presented concentration range of serially-diluted samples, including zero. The responses at equilibrium were fit (inset, red curve) against $\alpha^{E612K}$ concentrations with steady state affinity (SSA) binding model (Supplementary Methods, Equation 1) to obtain presented $K_{D1}$ values. The total number of residues on the ss template overhang of “0-X nt” is X+1 in the absence of a cognate nucleotide dGTP with 0 nt representing the first unpaired nucleotide at the p/t junction. Errors are standard errors of the fit.

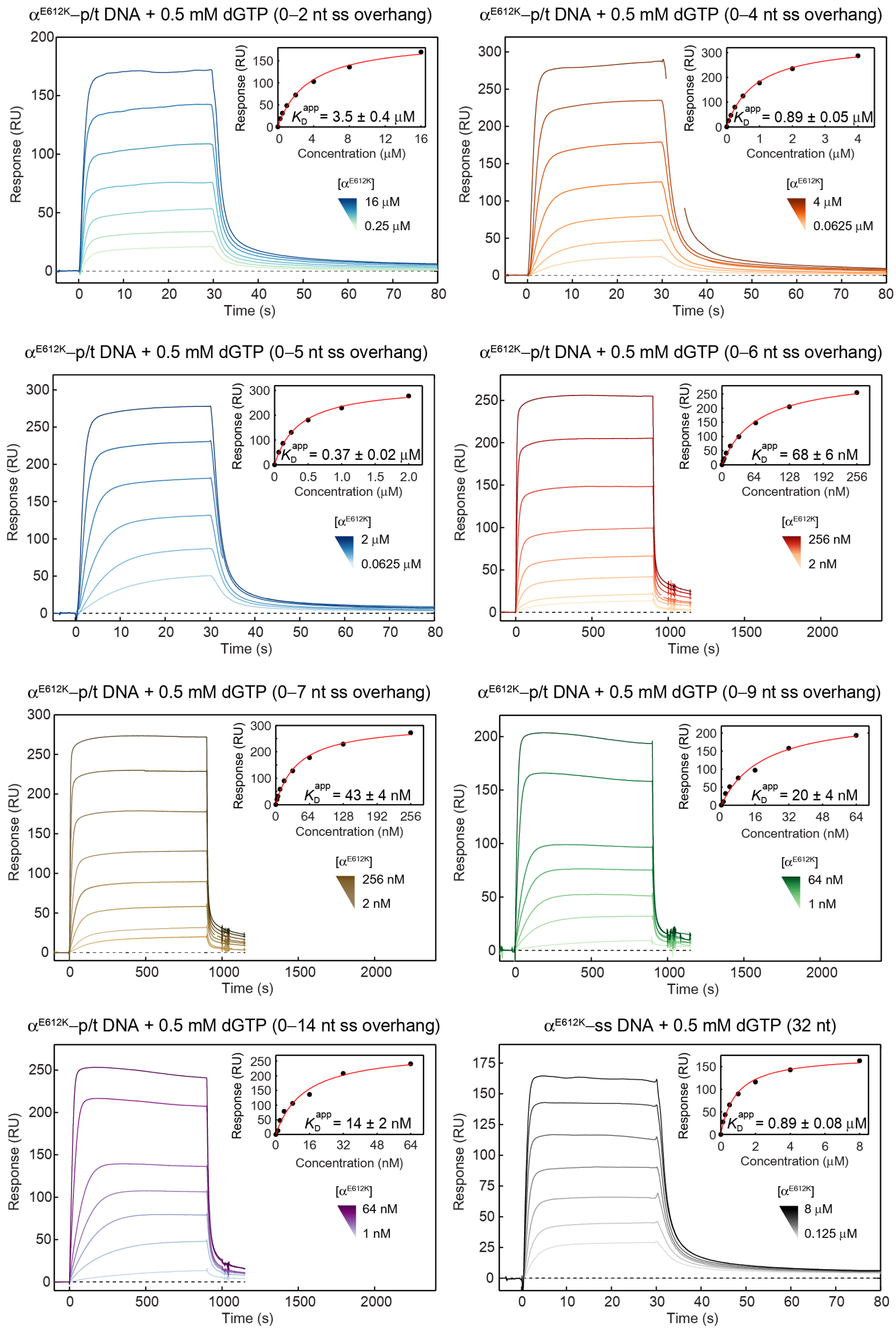

**Supplementary Figure 3. Binding of $\alpha^{E612K}$ to p/t DNA increases with increased number of residues on template overhang (for up to ~10 nt) and is ~15-fold stronger in the presence of saturating concentration of cognate nucleotide dGTP.** SPR sensorgrams showing association (30 s or 900 s) and dissociation of $\alpha^{E612K}$ to and from a p/t DNA or ss DNA over a presented concentration range of serially-diluted samples, including zero. Longer association times were necessary at lower concentrations to ensure signal saturation. The responses at equilibrium were fit (inset, red curve) against $\alpha^{E612K}$ concentrations with steady state affinity (SSA) binding model (Supplementary Data 1, Equation 8) to obtain presented apparent $K_D^{app}$ values. As seen with the 14 nt overhang, both $K_D$ and $K_D^{app}$ constants essentially reach their minimum values (240 ± 20 nM and 14 ± 2 nM, respectively), changing little compared to the 9 nt overhang (see also Supplementary Figure 2). Given that the calculated $K_{D2}^{app}$ (32 ± 8 μM; see Supplementary Data 1 and 2) is much (>10-fold) lower than the 0.5 mM dGTP concentration used in measurements, we conclude that the ternary $\alpha^{E612K}$p/t•DNA•dGTP complex is ~15-fold stronger than the binary $\alpha^{E612K}$p/t•DNA complex at saturating conditions. Residue 0 stands for the templating nucleotide that pairs with incoming dGTP, stabilizing the ternary complex. The total number of residues on the ss template overhang of “0-X nt” is then X. Errors are standard errors of the fit.

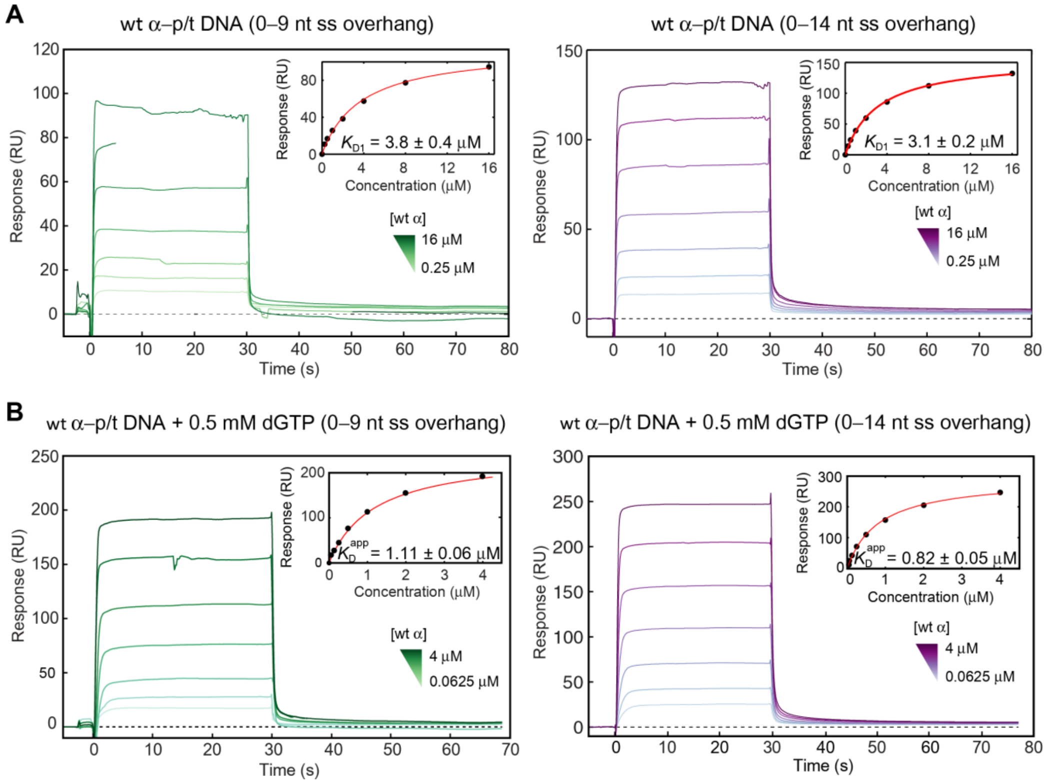

**Supplementary Figure 4. Binding of wt α to p/t DNA in the absence and presence of dGTP.** SPR sensorgrams show association (30 s) and dissociation of wt α from p/t DNAs without dGTP (A) and with cognate dGTP (B) over a presented concentration range of serially-diluted samples, including zero. The responses at equilibrium were fit (inset, red curve) against wt α concentrations with steady state affinity (SSA) binding model to obtain presented $K_{D1}$ (in case of binary complex formation) and apparent $K_D^{app}$ values (in case of mixed binary and ternary complex formation). Errors are standard errors of the fit.

Note that 0.5 mM dGTP stimulated binding of wt α to p/t DNAs with a 10 or 15 nt ss DNA region by 4-fold (comparing the formation of ternary vs binary complex), which is much less than the ~15-fold stimulation seen for binding of $\alpha^{E612K}$ to the same templates. This indicates that 0.5 mM dGTP must be a sub-saturating concentration for testing ternary complex formation of wt α with these templates, at least in the absence of the $\beta_2$ clamp and other factors in PolIII HE that normally stabilise complex formation in cell. Given these stabilizing factors, 0.5 mM dGTP would likely be a near saturating concentration. In support, the $K_m$ value for dNTP in the DNA chain elongation reaction was found to be 30 μM for the wild-type holoenzyme (Sugaya et al, 2002), a value > 10-fold lower than 0.5 mM (dGTP) used in our assay.

It is worth noting that usage of higher [dGTP] was unattainable since commercial nucleotide solutions have very high ionic strengths (> 1 M). When diluted to concentrations in the low mM range, dGTP still affected the ionic strength of injected solution, to which the α–DNA interaction is very sensitive. Likewise, weak binding of wt α to DNA prevented measurements of interactions with p/t DNA containing shorter DNA template overhangs due to the technical issues when very high α concentrations are used: a) limited solubility of highly concentrated α, and b) further incompatibility of glycerol needed for the stabilisation of even moderately concentrated α with SPR.

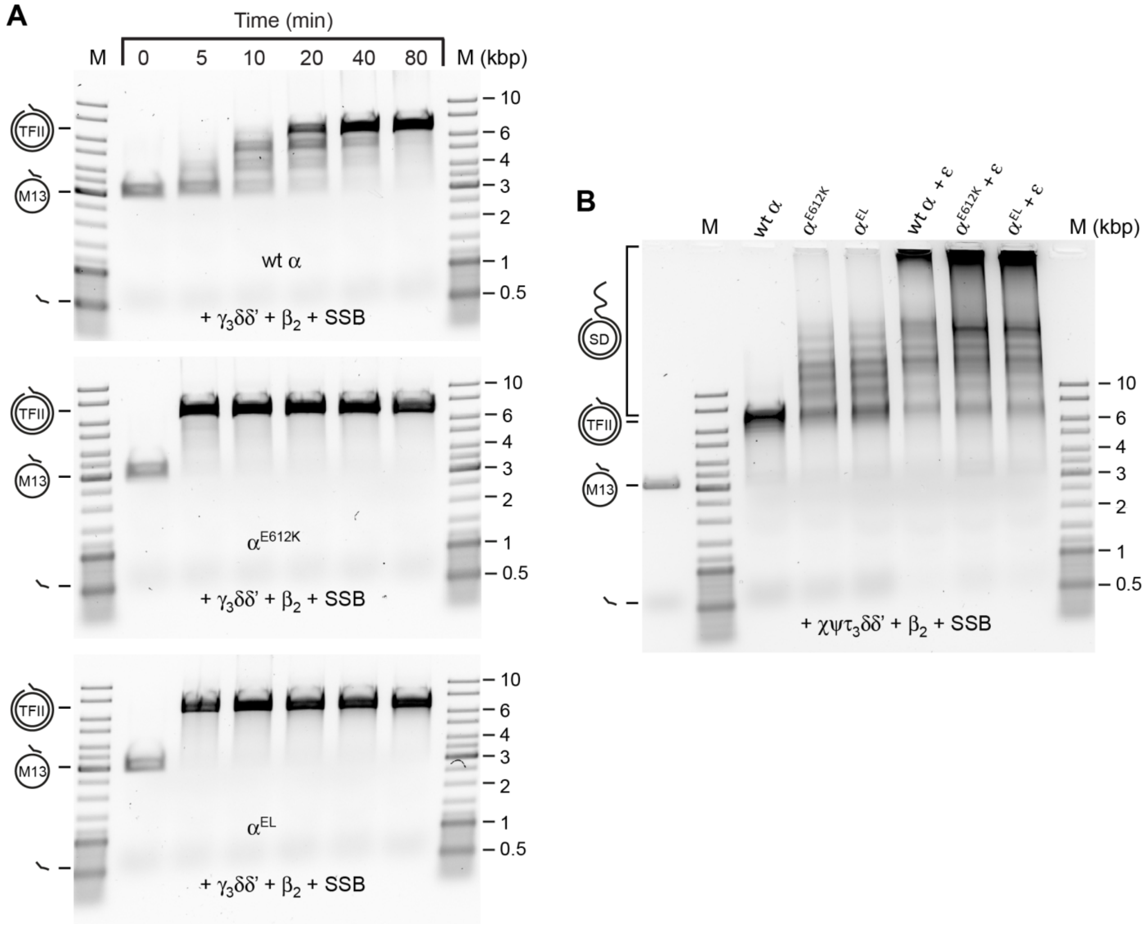

**Supplementary Figure 5. Both $\alpha^{E612K}$ and $\alpha^{EL}$ are more efficient polymerases compared to wt α.** (A) Time courses of primer extension "under difficult conditions" shows that while it took both $\alpha^{E612K}$ (middle panel) and $\alpha^{EL}$ (bottom panel) ≤ 5 min to produce unit length dsDNA (tailed form II, TFII) from flap-primed M13, wt α needed > 40 min to achieve the same result. Note, there are no products larger than TFII [strand displacement (SD) products] in the absence of τ. (B) Fixed time assays (20 min) of flap-primer extension on M13 ssDNA in the absence of ε subunit shows larger than unit length dsDNA products produced by SD synthesis of $\alpha^{E612K}$ and $\alpha^{EL}$ but not of wt α. In the presence of ε, however, all three polymerases were capable of strong SD synthesis, but more products are seen with $\alpha^{E612K}$ and $\alpha^{EL}$ relative to α. In both replication assays, no major difference in activities was observed between $\alpha^{E612K}$ and $\alpha^{EL}$.

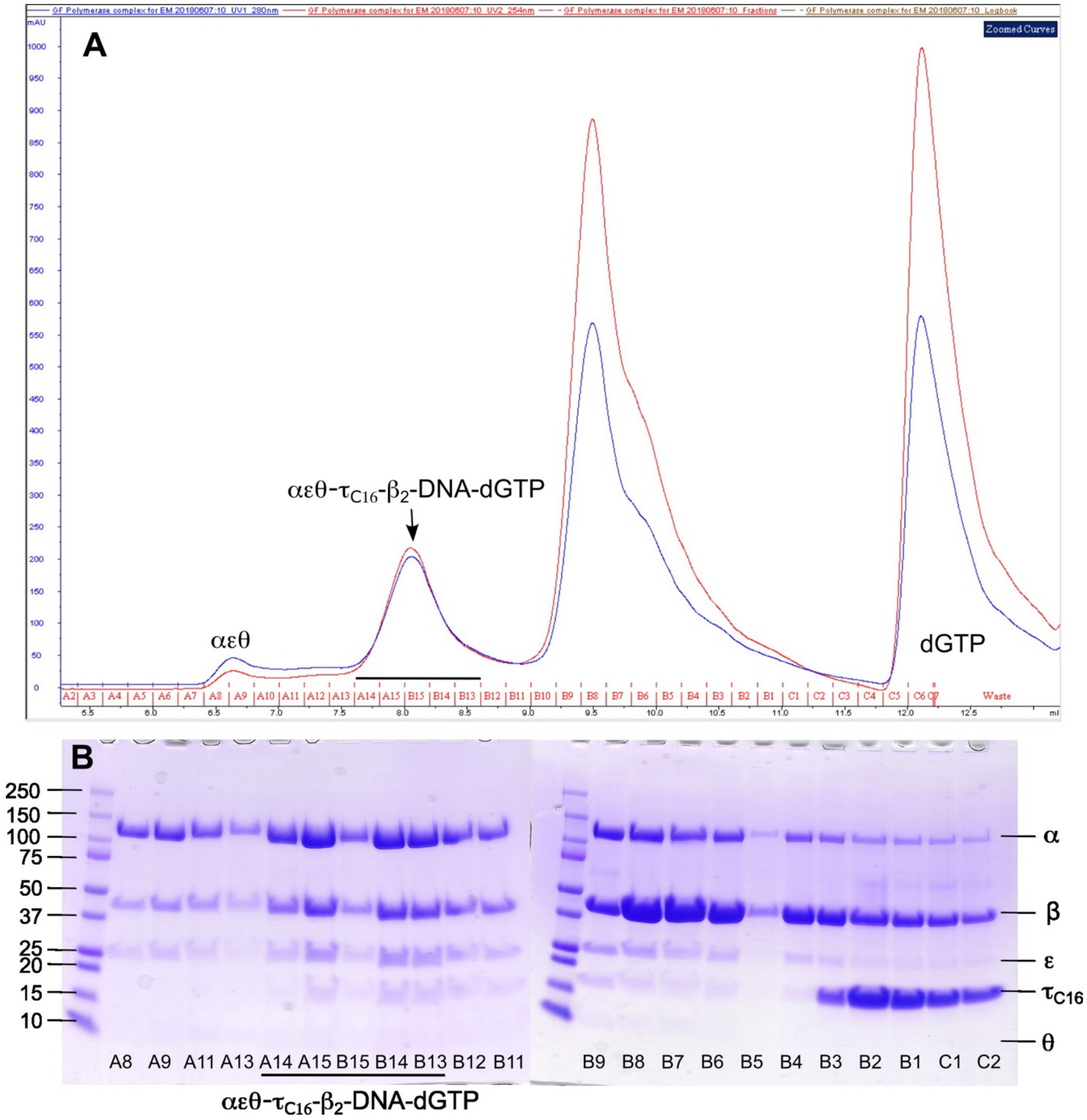

**Supplementary Figure 6. The $\alpha^{EL}\varepsilon^{DL}\theta\bullet\tau_{C16}\bullet\beta_2\bullet$DNA•dGTP complex was isolable by size exclusion chromatography.** (**A**) Chromatogram of the $\alpha^{EL}\varepsilon^{DL}\theta\bullet\tau_{C16}\bullet\beta_2\bullet$DNA complex on a 14-ml Wyatt WTC-030S5 SEC protein column. Blue curve is UV absorbance at 280 nm and red curve at 254 nm. The first peak (fractions A8–A10) contains "aggregated" $\alpha^{EL}\varepsilon^{DL}\theta$, which appears still able to interact with $\beta_2$. Concentrated $\alpha$ tends to form higher molecular weight complex. The second peak (A14–B13) contains the $\alpha^{EL}\varepsilon^{DL}\theta\bullet\tau_{C16}\bullet\beta_2\bullet$DNA complex, which is probably dGTP bound. The third peak (B9–C2) contains mixtures of $\alpha^{EL}\varepsilon^{DL}\theta\bullet\tau_{C16}$, excess $\beta_2$, $\tau_{C16}$ and DNA. The fourth contains the excess dGTP. (**B**) SDS-PAGE gels (4–20%) of individual fractions of the size exclusion chromatography. Note, repeated efforts to solve the structure of the $\alpha^{EL}\varepsilon^{DL}\theta\bullet\beta_2\bullet$DNA complex bound to a $\tau_{C16}$ had failed. Cryo-EM structures of the complex were later determined in the presence of either a full $\tau_3\delta\delta'\psi\chi$ clamp loader complex or $\tau_{C32}$ that contains both the helicase-interacting Domain IV and $\alpha$-binding Domain V with the same results.

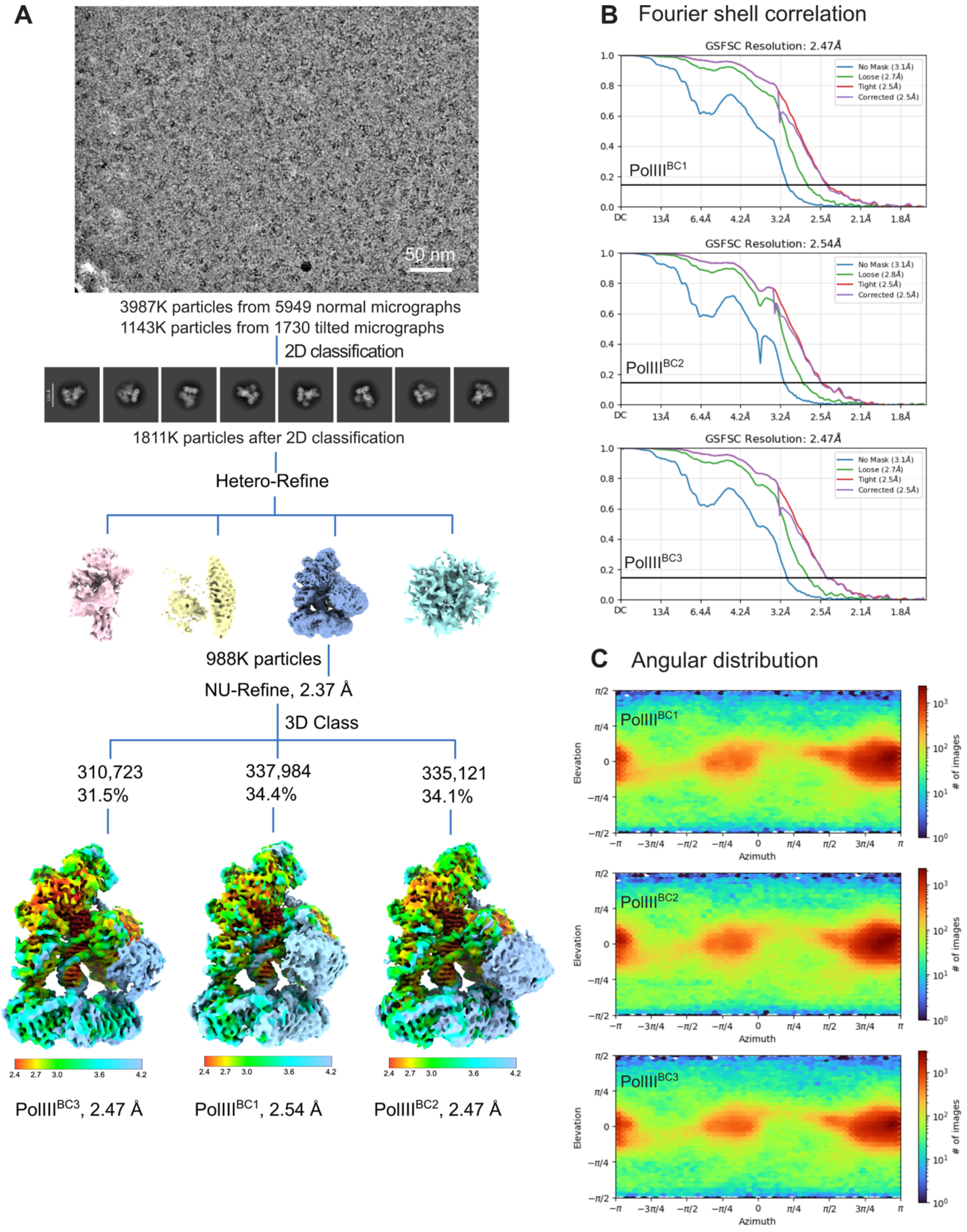

**Supplementary Figure 7. Processing of the cryo-EM data of the *E. coli* PolIII binary complexes (PolIII$^{BC}$), related to Figure 1.** (**A**), Cryo-EM image processing workflow. (**B**), Fourier shell correlation of the final density map. (**C**), Angular distribution plot of observed particle orientations.

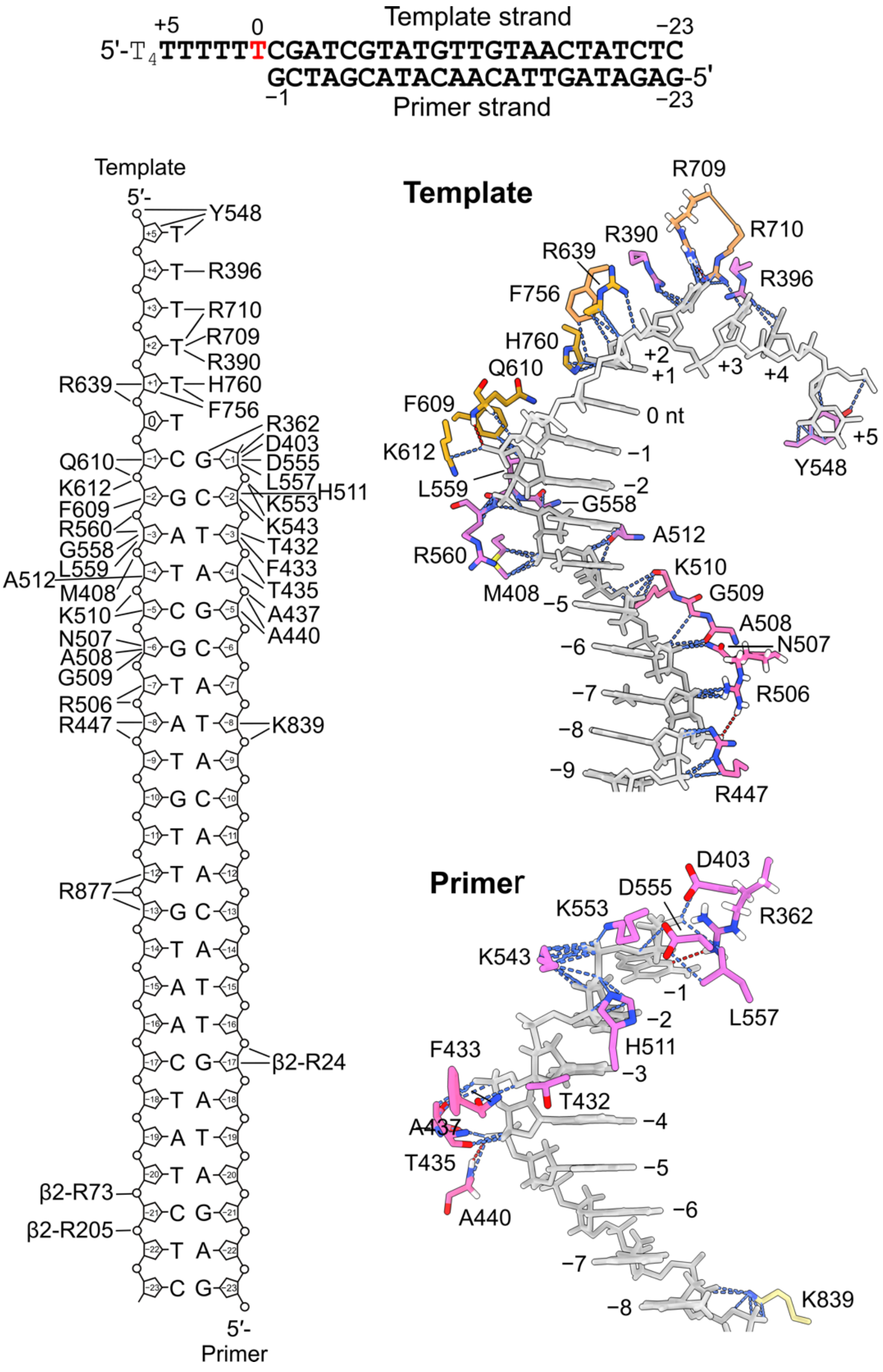

**Supplementary Figure 8. Protein–DNA interactions in the *E. coli* PolIII$^{BC1}$ binary complex, related to Figures 1 and 2.** Top, DNA sequences of the template and primer strands used in this experiment. DNA segments built in the structure are shown in bold. Residue positions relative to the "templating nucleotide" are shown above and below the sequences. 0 nt (red) on the template strand indicates the templating nucleotide for pairing with the incoming nucleotide (if present) during DNA polymerization. Bottom, schematic of protein–DNA interactions. DNA is shown as grey cartoons and DNA-interacting residues on $\alpha$ are shown as sticks. Residues on the palm, thumb, and index, middle, and ring fingers of $\alpha$ are coloured orchid, hot pink, sandy brown, goldenrod, and yellow, respectively. Only parts of DNA were shown (right). Protein–DNA contacts are shown as blue dashed lines and hydrogen bonds as red dashed lines. Hydrogens (white) on residues that hydrogen bond to DNA are also presented.

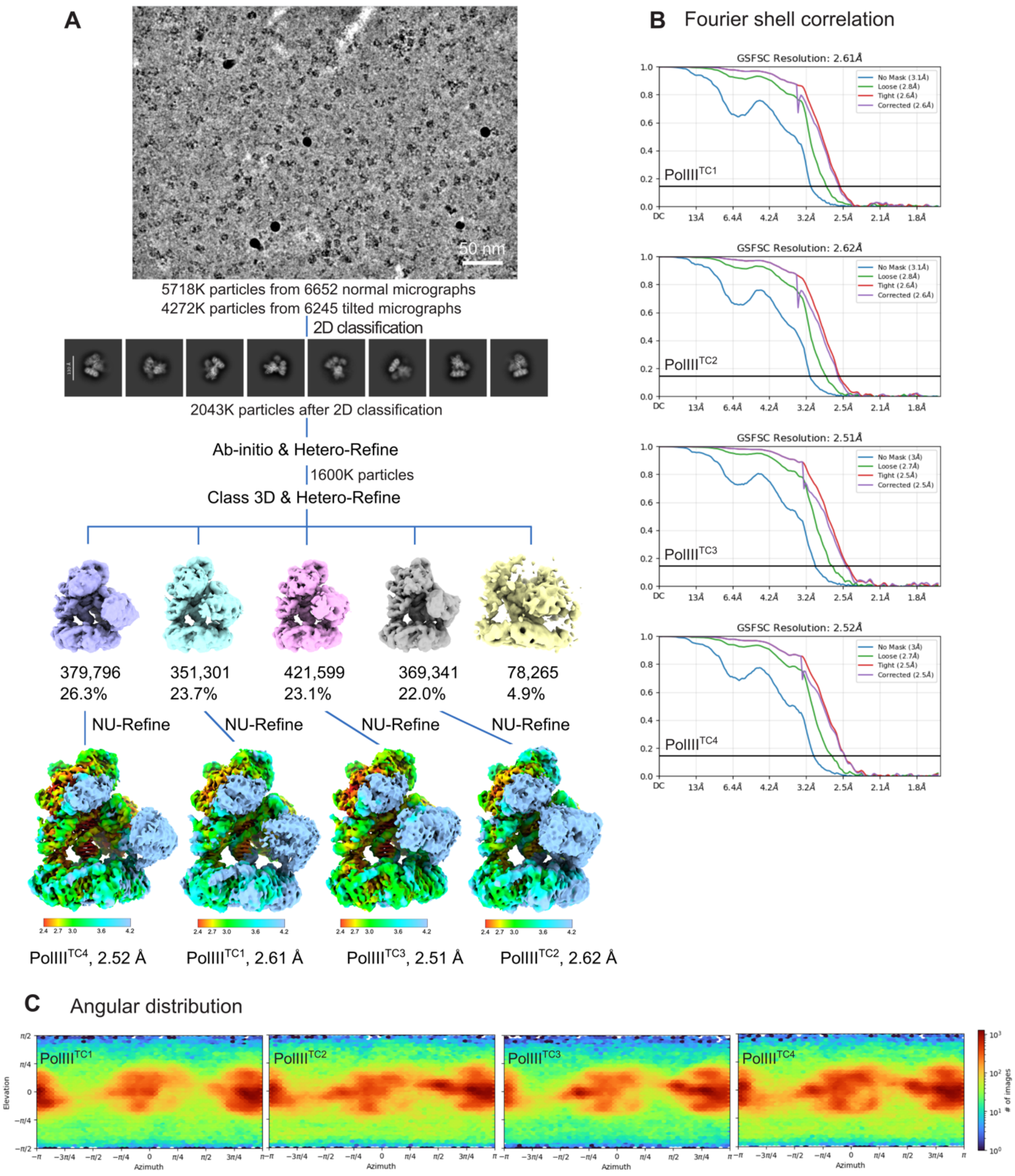

**Supplementary Figure 9. Processing of the cryo-EM data of the *E. coli* PolIII ternary complexes (PolIII$^{TC}$), related to Figure 3.** (**A**), Cryo-EM image processing workflow. (**B**), Fourier shell correlation of the final density map. (**C**), Angular distribution plot of observed particle orientations.

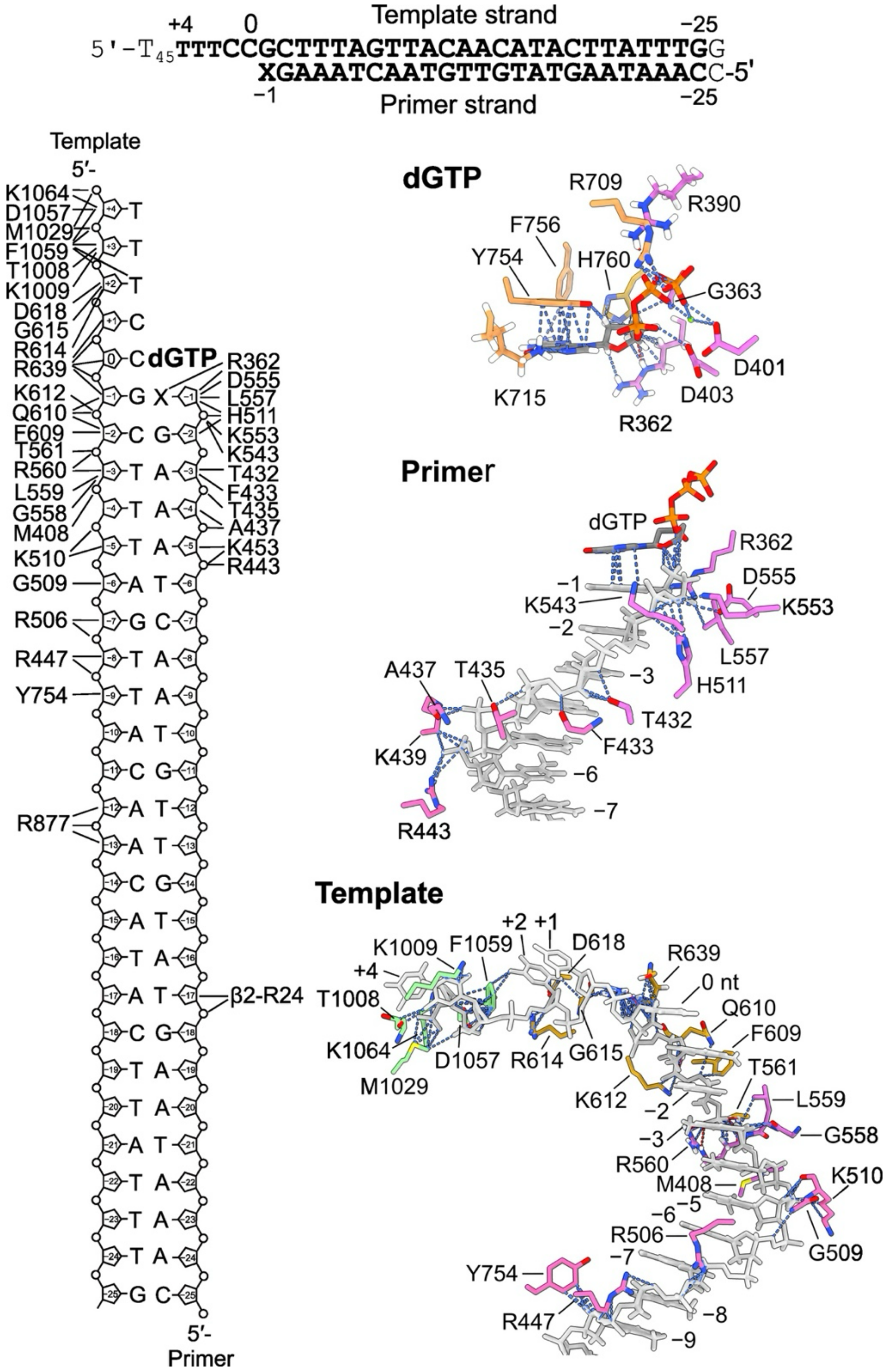

**Supplementary Figure 10. Protein–DNA interactions in the *E. coli* PolIII$^{TC2}$ ternary complex, related to Figures 3 and 4.** Top, DNA sequences of the template and primer strands used in this experiment. DNA segments built in the structure are shown in bold. 0 nt on the template strand indicates the templating nucleotide for pairing with the incoming nucleotide. Residue positions relative to the templating nucleotide are shown above and below the sequences. “X” denotes the 2′,3′-dideoxycytidine monophosphate (ddCMP) at the 3′-end of primer. Bottom, schematic of protein–DNA interactions. DNA is shown as grey cartoons and DNA-interacting residues on α are shown as sticks. Residues on the palm, thumb, and index, middle, and ring fingers of α are coloured orchid, hot pink, sandy brown, goldenrod, and yellow, respectively. Only parts of DNA were shown (right). Protein–DNA contacts are shown as blue dashed lines and hydrogen bonds as red dashed lines. Hydrogens (white) on residues that hydrogen bond to DNA are also presented.

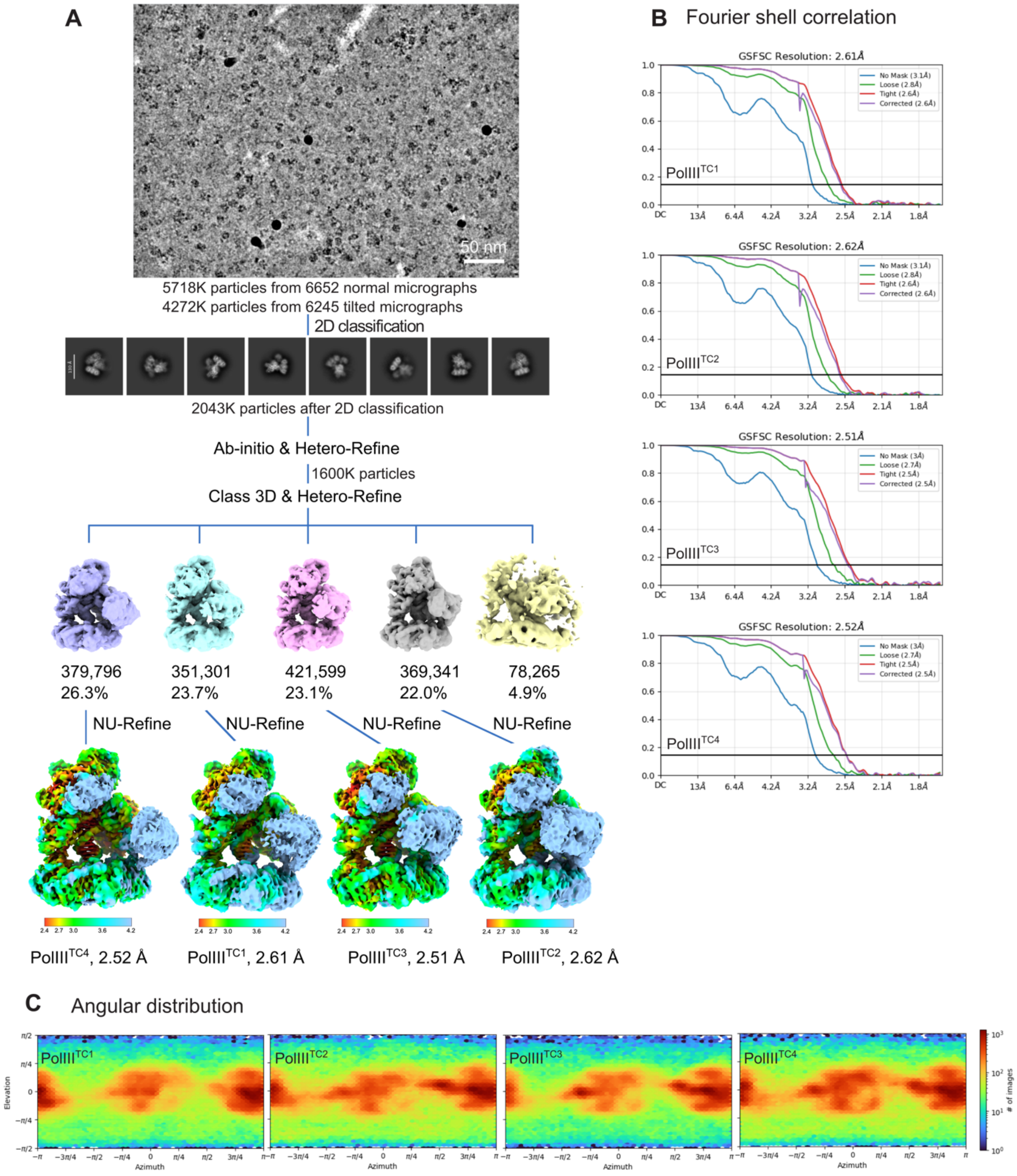

**Supplementary Figure 11. Processing of the cryo-EM data of the *E. coli* PolIII binary-like complexes (PolIII$^{BC*}$) and proofreading complexes (PolIII$^{PR}$) with mismatched p/t DNA, related to Figure 5.** (**A**), Cryo-EM image processing workflow. (**B**), Fourier shell correlation of the final density map. (**C**), Angular distribution plot of observed particle orientations.

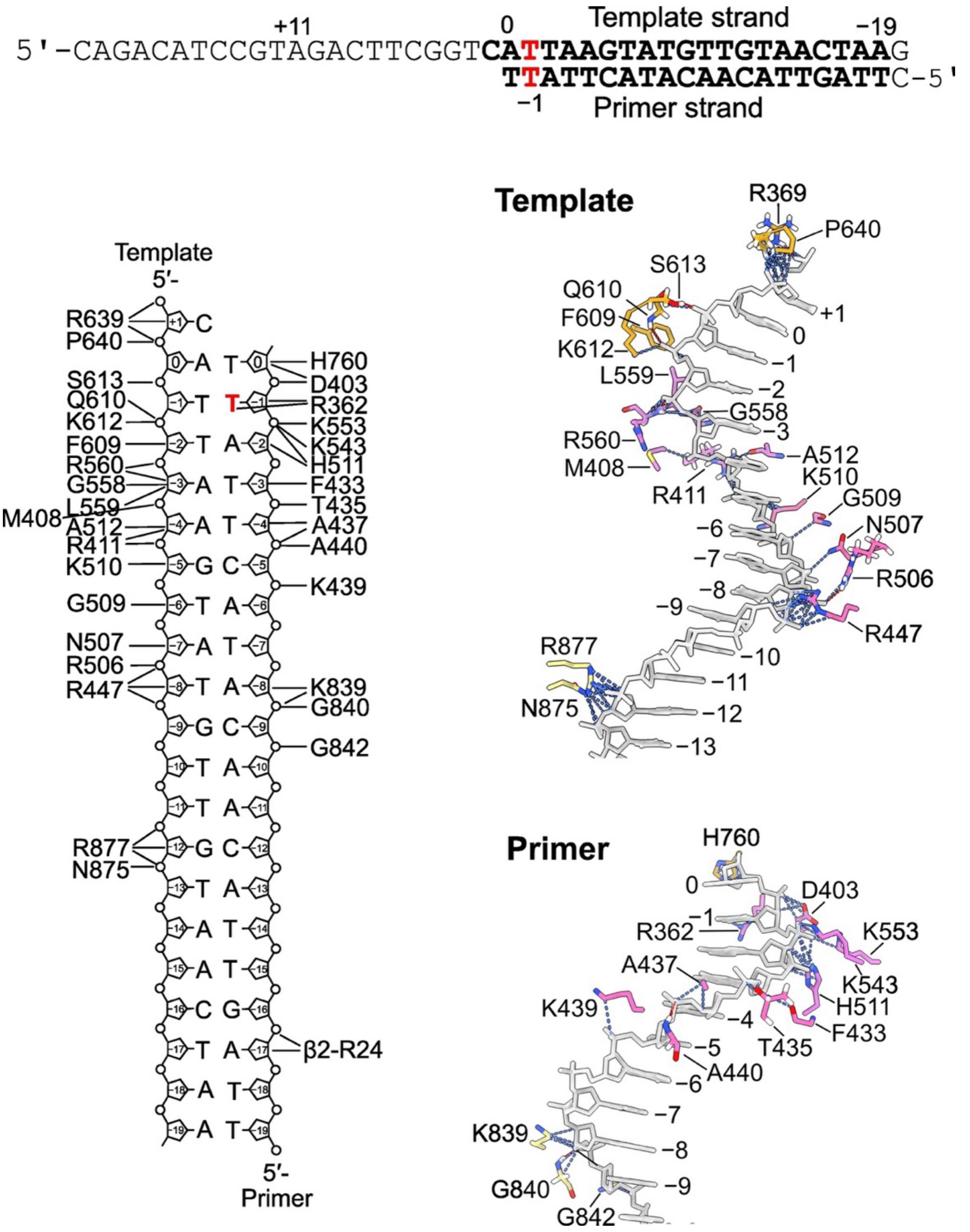

**Supplementary Figure 12. Protein–DNA interactions in the *E. coli* $\text{PolIII}^{\text{BC*1}}$ binary-like pre-proofreading complex, related to Figure 6**. Top, DNA sequences of the template and primer (nascent) strands used in this experiment. DNA segments built in the structure are shown in bold. 0 nt denotes the nucleotides in the active site of $\alpha$ had DNA not retracted for proofreading, *i.e.*, the templating nucleotide on the template strand and newly inserted nucleotide on the primer strand (see Figure 6A), while −1 nt (red) on the primer strand is the mismatched nucleotide. Residue positions relative to the "0 nt templating nucleotide" are shown above and below the sequences. Unlike $\text{PolIII}^{\text{BC*}}$, the 3′-end of the primer strand in the binary-like $\text{PolIII}^{\text{BC*1}}$ complex sits in the active site of $\alpha$. Bottom, schematic of protein–DNA interactions. DNA is shown as grey cartoons and DNA-interacting residues on $\alpha$ are shown as sticks. Residues on the palm, thumb, and index, middle, and ring fingers of $\alpha$ are coloured orchid, hot pink, sandy brown, goldenrod, and yellow, respectively; and those on $\varepsilon$ medium purple. Protein–DNA contacts are shown as blue dashed lines and hydrogen bonds as red dashed lines. Hydrogens (white) on residues that hydrogen bond to DNA are also presented.

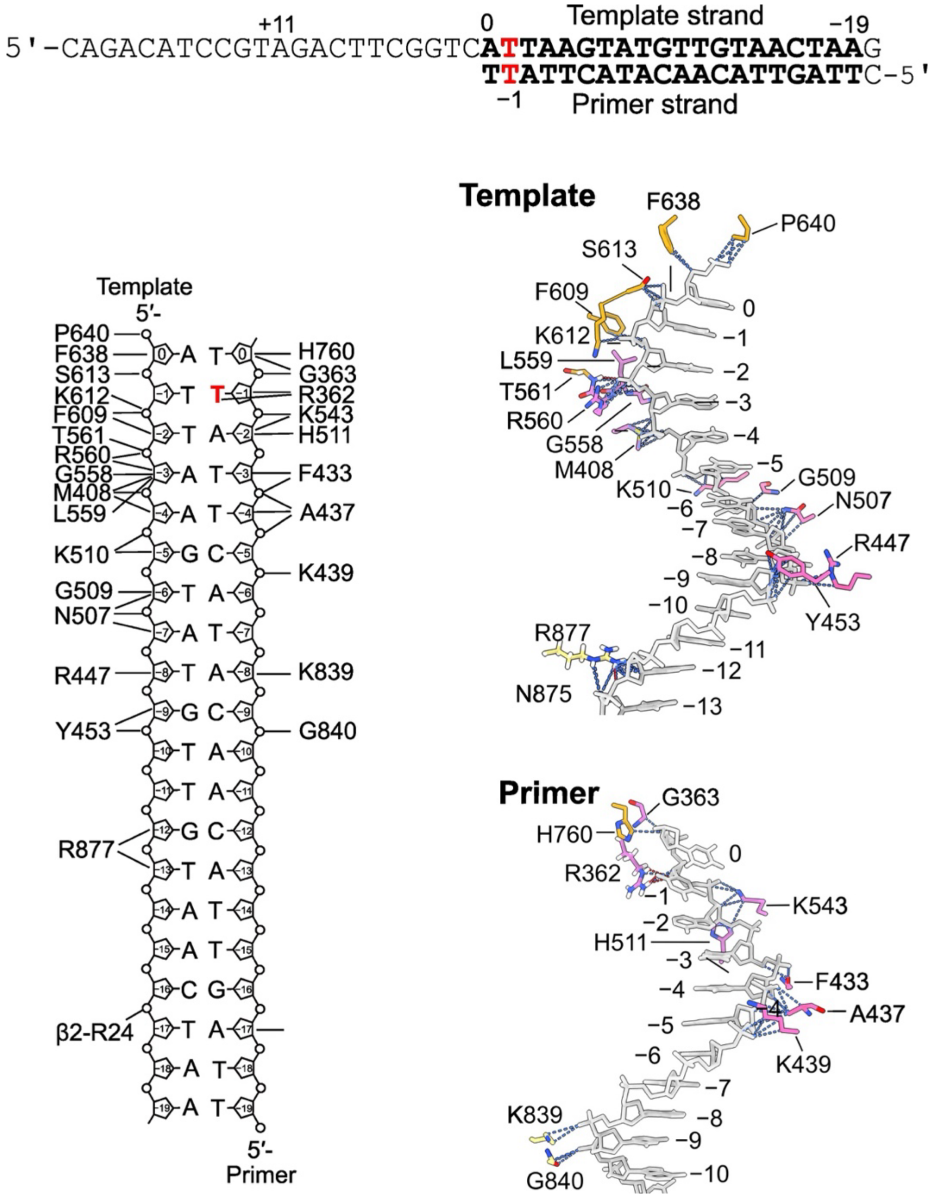

**Supplementary Figure 13. Protein–DNA interactions in the *E. coli* PolIII$^{BC*2}$ binary-like pre-proofreading complex, related to Figure 6**. Top, DNA sequences of the template and primer (nascent) strands used in this experiment. DNA segments built in the structure are shown in bold. 0 nt denotes the nucleotides in the active site of $\alpha$ had DNA not retracted for proofreading, *i.e.*, the templating nucleotide on the template strand and newly inserted nucleotide on the primer strand (see Figure 6A), while −1 nt (red) on the primer strand is the mismatched nucleotide. Residue positions relative to the "0 nt templating nucleotide" are shown above and below the sequences. Unlike PolIII$^{BC*}$, the 3′-end of the primer strand in the binary-like PolIII$^{BC*1}$ complex sits in the active site of $\alpha$. Bottom, schematic of protein–DNA interactions. DNA is presented as grey cartoons and DNA-interacting residues on $\alpha$ are shown as sticks. Residues on the palm, thumb, and index, middle, and ring fingers of $\alpha$ are coloured orchid, hot pink, sandy brown, goldenrod, and yellow, respectively; and those on $\varepsilon$ medium purple. Protein–DNA contacts are shown as blue dashed lines and hydrogen bonds as red dashed lines. Hydrogens (white) on residues that hydrogen bond to DNA are also presented. The protein–DNA interactions in PolIII$^{BC*2}$ appear less intensive than in PolIII$^{BC*1}$. Note, the DNA density in PolIII$^{BC*2}$ is much weaker than that in PolIII$^{BC*1}$, therefore, the modelling can be less accurate. However, the weaker density itself is an indication of more flexibility and weaker protein–DNA interactions.

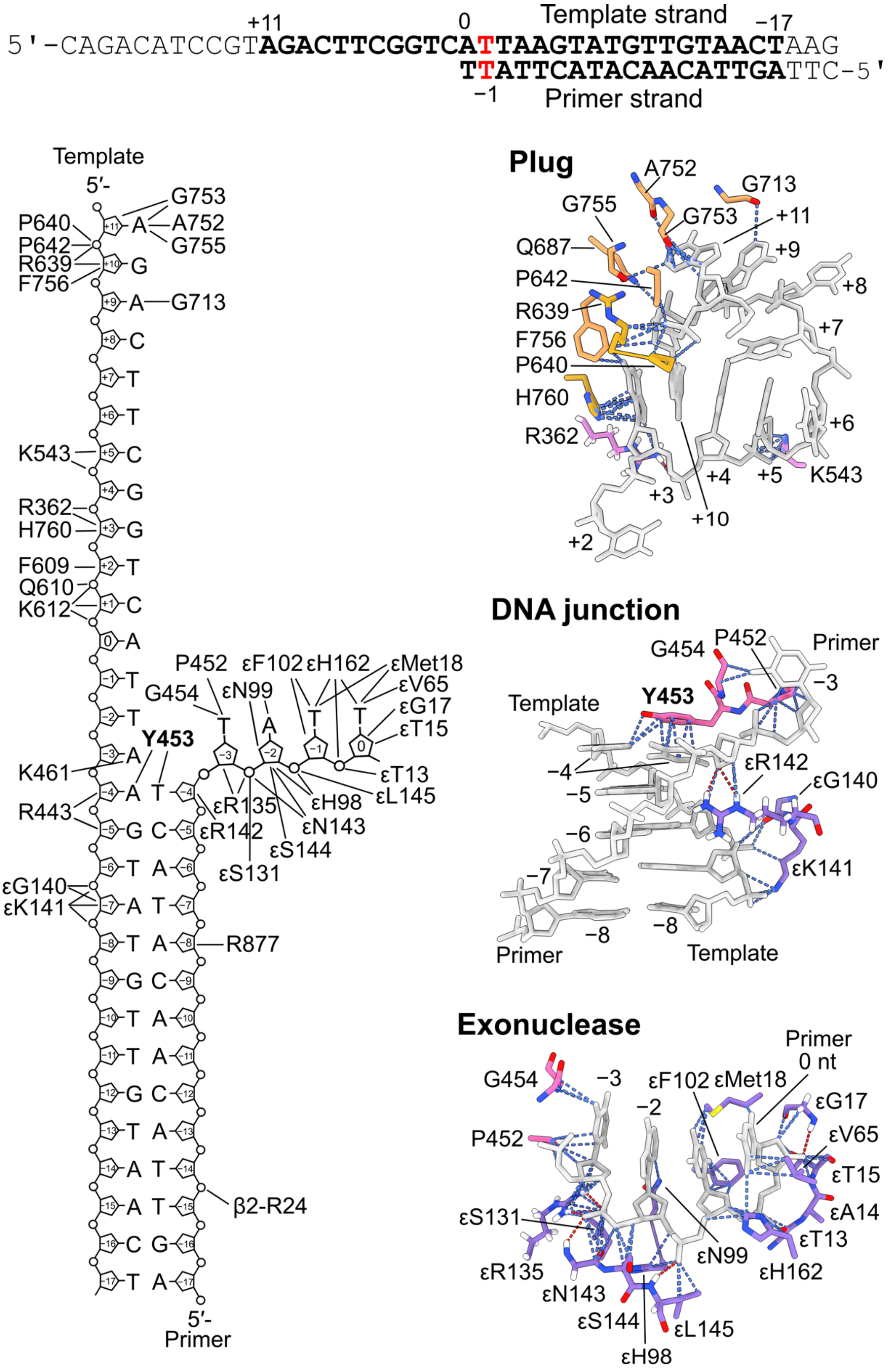

**Supplementary Figure 14. Protein–DNA interactions in the *E. coli* PolIII[PR1] proofreading complex, related to Figure 6**. Top, DNA sequences of the template and primer (nascent) strands used in this experiment. DNA segments built in the structure are shown in bold. 0 nt denotes the nucleotides in the active site of α had DNA not retracted for proofreading, *i.e.*, the templating nucleotide on the template strand and newly inserted nucleotide on the primer strand (see Figure 6A), while −1 nt (red) on the primer strand is the mismatched nucleotide. Residue positions relative to the

“0 nt templating nucleotide” are shown above and below the sequences. Unlike the binary-like $PolIII^{BC*}$ complexes, the primer strand in the $PolIII^{PR1}$ proofreading complex retracted from the polymerase’s active site and reached to the exonuclease active site, while the template overhang formed a plug-like structure anchored in the centre channel of $\alpha$. Bottom, schematic of protein–DNA interactions in the proofreading complex. DNA is presented as grey cartoons and DNA-interacting residues on $\alpha$ or $\varepsilon$ as sticks. Residues on the palm, thumb, and index, middle, and ring fingers of $\alpha$ are coloured orchid, hot pink, sandy brown, goldenrod, and yellow, respectively; and those on $\varepsilon$ medium purple. Protein–DNA contacts are shown as blue dashed lines and hydrogen bonds as red dashed lines. Hydrogens (white) on residues that hydrogen bond to DNA are also presented. Note, Lys612 interacts with −2 nt of the template strand in the PolIII binary/ternary complexes while it contacts +1 nt in the proofreading complex, illustrating the extent of DNA retraction relative to $\alpha$.

**Captions of Supplementary movies**

**Supplementary movie 1. Conformational changes of the palm and thumb of $\alpha$ on DNA binding.** The movie shows how the palm and thumb of $\alpha$ undergo conformational changes to adapt to DNA binding. The P$\beta$5 strand and the following helix on the $\beta_2\alpha$ structural motif on the palm are coloured orchid and the rest of the palm white. The antiparallel $\beta$-strands on the thumb are coloured hot pink and the rest light grey. The other domains of $\alpha$ are omitted for clarity. Residues involved in DNA interactions are shown as sticks.

**Supplementary movie 2. Conformational differences of the four PolIII ternary complexes.** Morphing of the four $PolIII^{TC}$ structures, from $PolIII^{TC1}$ to $PolIII^{TC4}$, showing the differences of these structures.

**Supplementary movie 3. The template overhang did not swing out of the central channel of $\alpha$ spontaneously**. The movie shows the trajectory of the 1-$\mu$s molecular dynamics simulation of $PolIII^{BC}$. The ss template overhang does not swing out of the central channel of $\alpha$ spontaneously due to electrostatic interactions with polar and positively charged residues on the surface of the channel. Red spheres denote phosphate atoms on the template overhang, blue spheres represent positively charged residues on the surface of the channel that interact with the overhang.

**Supplementary movie 4. Binding of dGTP expels the template overhang out of the central channel of $\alpha$**. The movie shows the trajectory of the 1-$\mu$s molecular dynamics simulation of $PolIII^{BC}$ with a dGTP placed at the nucleotide insertion site, showing how binding of a dGTP (spheres) expels the template overhang (in rainbow colours) out of the central channel of $\alpha$, which transiently interacts with OB.

**Supplementary Methods**

**Primer extension under difficult conditions**

Assays (A) in the absence of exonuclease subunit $\varepsilon(\theta)$ and $\tau$ contained 2.8 nM 5'-flap-primed M13 (6.4 kb) DNA template (Jergic et al., 2013), 1 mM ATP, 0.42 mM of each dNTP, 10 mM dithiothreitol, 20 nM $\gamma_3\delta\delta'$ clamp loader, 30 nM $\beta_2$, 600 nM SSB and 90 nM of either wt $\boldsymbol{\alpha}$, $\boldsymbol{\alpha}^{E612K}$ or $\boldsymbol{\alpha}^{EL}$ in 20 mM Tris.HCl pH 7.6, 100 mM NaCl, and 8 mM $MgCl_2$, in final volume of 12 µl.

Replication components (except DNA) were mixed and treated for 5 min at room temperature to allow proteins to interact, then cooled on ice and DNA added. The reactions were initiated by quick transfer to a 30˚C water bath, and quenched at indicated times by addition of 11 µl quenching buffer (200 mM EDTA pH 8.0, 0.08% (w/v) bromophenol blue, 0.08% (w/v) xylene cyanol, 10% (v/v) glycerol and 2% (w/v) SDS). Reaction mixtures were treated for 2 min at 42˚C, loaded onto a 0.7% agarose gel in 2×TBE buffer and electrophoresis carried out at 80 V for 130 min. Gels were stained for 75 min in 200 ml of SYBR® gold nucleic acid stain at the concentration suggested by the supplier (Invitrogen, Carlsbad, CA). The DNA products were visualized using a UV transilluminator. A sample of GeneRuler™ 1 kb Plus DNA Ladder (Thermo Fisher Scientific) was loaded in two lanes of each gel, as a reference.

**Strand displacement assay**

Assay (B). The coupled Pol III primer extension–strand displacement (SD) rolling circle assay in the absence of $\varepsilon(\theta)$ contained 2.6 nM 5'-flap-primed M13 ssDNA template, 1 mM ATP, 0.42 mM of each dNTP, 25 nM $\chi\psi\tau_3\delta\delta'$, 240 nM $\beta_2$, 750 nM SSB, and 90 nM of either wt $\boldsymbol{\alpha}$, $\boldsymbol{\alpha}^{E612K}$ or $\boldsymbol{\alpha}^{EL}$, in 20 mM Tris.HCl pH 7.6, 70 mM NaCl, and 8 mM $MgCl_2$, in final volume of 13.5 µl. When present, $\varepsilon$ was added to reactions at 200 nM. Replication components (except DNA) were mixed and treated for 5 min at room temperature, then cooled on ice and DNA added. The reactions were initiated by transfer to a 30°C water bath, and quenched after 20 min by addition to 12 µl of quenching buffer. Reaction mixtures were then treated for 5 min at 42˚C, loaded onto a 0.66% agarose gel in 2×TAE buffer and the DNA products resolved by electrophoresis at 80 V for 130 min. Gel was stained for 75 min in 200 ml of SYBR® gold nucleic acid stain at the concentration suggested by the supplier (Invitrogen, Carlsbad, CA). The DNA products were visualized using a UV transilluminator. A corresponding DNA template sample was loaded in one lane as reference, as well as a sample of GeneRuler™ 1 kb Plus DNA Ladder (Thermo Fisher Scientific).

**Polymerase–DNA interaction analysis by SPR**

Interactions between $\boldsymbol{\alpha}^{E612K}$ and wt $\alpha$ with various primed-template (p/t) DNAs that differ in number of residues in the ss DNA region were measured in the absence and presence of cognate nucleotide dGTP. All hybridizations and measurements were performed at 25°C in SPR buffer: 30 mM Tris, pH 7.6, 50 mM NaCl, 12 mM $MgCl_2$, 0.5 mM EDTA, 0.5 mM DTT containing 0.005% surfactant P20.

First, a streptavidin coated (SA) chip was activated with three sequential injections of 1M NaCl, 50 mM NaOH at 5 µl/min for 60 s. The surface was further stabilized by two injections of 1M $MgCl_2$, with the same contact times and flow rates. Then biotinylated primer 5'-bio-AAAACGAAAAATAAGTATGTTGTAACTAAAG(ddC)-3' (32-mer) was immobilized onto the surface. This primer was used as a template when binding to ss DNA was considered. To assemble a particular p/t DNA, an appropriate template strand from the partial set of oligos 5'-$(C)_{x=3-15}$GCTTTAGTTACAACATACTTATTTTTCGTTTT-3' (500 nM) was hybridized on-line by injecting it at 5 µl/min for 300 s. Note that we adopted such nomenclature for assembled p/t DNA where the first dC at the p/t junction is labelled as 0 nt. This nucleotide is also termed templating nucleotide given it tends to pair with cognate dGTP when it is present in the reaction.

Interactions between $\boldsymbol{\alpha}^{E612K}$ with p/t DNA in the absence of a cognate nucleotide dGTP were carried out by sequential coinjections of one or two appropriate concentration series in SPR buffer (zero and presented solutions of serially diluted samples) at 20 μl/min for 30 s, followed by dissociation in the same buffer over 60 s. In the presence of dGTP (0.5 mM), some of $\boldsymbol{\alpha}^{E612K}$ were measured as described above, while in case of stronger interactions of $\boldsymbol{\alpha}^{E612K}$ with p/t DNAs containing longer templates, coinjections were performed at 15 μl/min for 900 s, followed by dissociation in the same buffer over 100 s. Longer injection times were necessary for the signal to reach equilibrium (saturation) at lower utilised concentrations of $\boldsymbol{\alpha}^{E612K}$ that were necessary for optimal curve fitting. All measurements involving wt α, with and without dGTP, were performed as above for samples of $\boldsymbol{\alpha}^{E612K}$ that did not contain dGTP, except that the flow rate was 100 μl/min. No dGTP was present in the dissociation phase, which is inconsequential given that studies are based on measured responses at equilibrium ($R_{eq}$) upon binding of polymerase. Likewise, inconsequential were sporadic missing (deleted) pieces of sensorgrams during dissociation due to protruding spikes of air.

Between successive injections within one concentration series, surfaces (p/t DNAs) were regenerated with 1 min injections of 1 M $MgCl_2$ at 5 μl/min. When a new (different) p/t DNA was to be assembled in place of an old, the template strand from previous concentration run was removed with two 1 min injections of 1M NaCl, 50 mM NaOH at 5 μl/min, and a new template strand hybridized as previously described.

The final sensorgrams were subtracted from an unmodified ligand flow path using BIAcore software and then zero subtracted using Mathlab. Equilibrium binding parameters (dissociation constants, $K_D$) were determined using a 1:1 steady state affinity (SSA) binding model:

$$R_{\text{eq}} = R_{\max}\left(\frac{[A]}{[A]+K_{\text{D}}}\right) \qquad \text{(Equation 1)}$$

where $R_{max}$ corresponds to the response when all the immobilized ligands (DNA templates) on the surface are saturated with the analyte A (polymerase), $K_D$ is the dissociation constant, and [A] is the concentration of analyte in solution for which $R_{eq}$ represents the corresponding equilibrium binding response upon interaction with surface immobilized DNA. Likewise, apparent dissociation constants $K_D^{app}$ obtained in the presence of complementary nucleotide dGTP were determined using the similar SSA binding model (Equation 8) that is fully derived in Supplementary Data 1.

#### Expression trials of the $\alpha^{Y754A}$ and $\alpha^{Y754F}$ variants

Plasmids encoding $\alpha^{Y754A}$ or $\alpha^{Y754F}$ were transformed into both *E. coli* BL21-AI and BL21 (λDE3)/pLysS stains by heat shock. Eight single colonies from each transformation were picked for expression trials. Trials either started by inoculating cells scraped from agar plates into 6 ml of LB to $OD_{600}$ ~0.2, or by adding 120 μl of overnight liquid LB cultures into 6 ml of LB. The cells were grown at 37°C until $OD_{600}$ reached ~0.6. 1 mM of IPTG was then added to induce protein expression. For BL21-AI cells, 0.2% of L-(+)-arabinose was also added. The cells were shaken at 30°C for 4 h and samples were run on SDS-PAGE gels to check for protein expression. No α was seen in three separate trials.

#### Size exclusion chromatography of the αεθ•$\tau_{C16}$•$\beta_2$•DNA•dGTP complex

The αεθ•$\tau_{C16}$•$\beta_2$•DNA sample for gel filtration was prepared by mixing 50 μl of 52.3 μM $\alpha^{EL}\varepsilon^{DL}\theta$ with 18.8 μl of 200 μM $\tau_{C16}$, 26 μl of 130 μM $\beta_2$, and 18 μl of 150 μM p/t DNA. 1 μl of 1M dithiothreitol and 2 μl of 200 mM $MgCl_2$ were then added and the mixture was incubated on ice for 20 minutes before loading onto a 14-ml Wyatt WTC-030S5 SEC protein column equilibrated with 30 mM Tris-HCl, pH 7.6, 5 mM $MgCl_2$, 3 mM dithiothreitol, 0.25 mM EDTA, 70 mM potassium glutamate and 100 μM dGTP. Chromatography was performed at 4˚C with a flow rate of 0.5 ml/min. Elution of the

$\alpha\epsilon\theta \bullet \beta_2 \bullet \tau_{C16} \bullet$DNA•dGTP complex was complete within 18 min. Fractions of 200 μl were collected. DNA was prepared by annealing Oligo 3 [5′-CCAAATAAGTATGTTGTAACTAAAG(ddC)] (TriLink Biotechnologies, PAGE purified) and Oligo 4 (5′-$T_{48}$-CCGCTTTAGTTACAACATACTTATTTGG) (IDT).

**Molecular dynamics simulations**

To prepare the initial structures for simulation, the DNA sequences of the PolIII binary complex were changed to be the same as that of ternary complex. The ss DNA regions of the template overhangs in both binary and ternary complexes were then extended to 10 nt by adding extra dTs and the ds DNAs were lengthened by 4 base pairs. The 3'-ddCMP on the primer strand of the ternary complex was changed to dCMP. So, the final sequences of the template strands become: 5′-TTTTTTTTCCGCTTTAGTTACAACATACTTATTTGCGCG; and the primer strands: 5′-CGCGCAAATAAGTATGTTGTAACTAAAGC. For simulation of PolIII$^{BC}$ with dGTP, a dGTP was manually positioned in the active site of PolIII$^{BC}$ like the dGTP in PolIII$^{TC}$. The complexes were subsequently immersed in a rectangular box, with a 14 Å buffering between the box boundaries and the outmost atoms of the complexes. To neutralize the system, NaCl was added to 50 mM and $MgCl_2$ to 10 mM. The ff14SB (Maier, 2015) and OL24 (Zgarbová, 2025) force fields were employed for protein and DNA, respectively. Water was treated using SPC/E model (Berendsen,1987) and parameters of dGTP derived from the GTP force field (Meagher, 2003).

The entire systems were first minimized for 10,000 steps with all the heavy atoms restrained (force constant 5 kcal/mol/Å$^2$), except for water and ions. The initial 5,000 steps were conducted using steepest descent algorithms, followed by conjugate gradient for 5,000 steps. Temperature was then heated to 310 K within 50 ps under the canonical ensemble (NVT) with restraints maintained. Followed by 10 ns isothermal-isobaric ensemble (NPT), the densities were equilibrated to an optimized value with the restrained force constant evenly decreased from 5 to 1 kcal/mol/Å$^2$. All restraints were removed during production and whole simulations last for 1000 ns under NPT. The temperature and pressure were maintained at 310 K and 1 atm using Langevin dynamics with a collision frequency of 3 ps$^{-1}$ and isotropic position scaling, respectively.

All molecular dynamics (MD) simulations were conducted under the periodic boundary conditions with Amber 22 package suite (Case, 2022). The long-range electrostatic interactions were treated by the particle mesh Ewald method (PME) (Darden, 1993). For both the short-range electrostatic and van der Waals (vdW) interactions, a cut-off of 8 Å was applied. The integration time step was extended to 4 fs with hydrogen mass repartitioning (Hopkins, 2015). Bonds involving hydrogen were constrained using the SHAKE algorithm (Ryckaert, 1977). All the analysis performed here were done leveraging the CPPTRAJ module of Ambertools (Case, 2023), along with our homemade scripts embedded with VMD 1.9.4 (Hemphrey, 1996). All parameters and scripts used for preparing and analysing MD simulations have been deposited at https://github.com/QZResearch/PolIII_ZQ_Xu/tree/main.

**Supplementary Data**

**Supplementary Data 1: Model for fitting the data in SPR studies obtained in the presence of complementary nucleotide dGTP**

We adapted the following, most simplistic, general working model for the formation of binary and ternary complexes on the SPR chip surface in our analysis:

$$\alpha + \mathrm{DNA} \rightleftarrows \alpha.\mathrm{DNA} + \mathrm{dGTP} \rightleftarrows \alpha.\mathrm{DNA.dGTP}$$

where $\frac{k_2}{k_1} = K_{\mathrm{D1}}$ is a dissociation constant for the reaction $\mathrm{DNA} \rightleftarrows \alpha + \mathrm{DNA}$. It thus reports on the stability of the binary complex. This value is independently obtained from the SPR measurements of α binding to immobilized DNA, in the absence of dGTP. This interaction represents a state when the ss region of p/t DNA is bound in the central cavity of α, since this is the only binary state we could observe.

The apparent dissociation constant $K_{\mathrm{D}}^{\mathrm{app}}$ derived within the further text (Equation 8) reports on the increased stabilisation of the α–DNA interaction in the presence of a cognate nucleotide (dGTP); in particular its smaller value (reduction) relative to $K_{\mathrm{D1}}$ value obtained for the same p/t DNA informs on the increased efficiency of formation of α.DNA.dGTP ternary complexes at utilised experimental conditions.

Finally, $\frac{k_{4*}}{k_{3*}} = K_{\mathrm{D2}}^{\mathrm{app}}$ for the reaction $\alpha.\mathrm{DNA.dGTP} \rightleftarrows \alpha.\mathrm{DNA} + \mathrm{dGTP}$. We defined it as an apparent constant because it also incorporates (not presented) a simplified hypothesized non-dimensional linear term stemming from the conformational change(s) in α.DNA upon nucleotide binding, as seen in the structure. $K_{\mathrm{D2}}^{\mathrm{app}}$ can be calculated from Equation 10, once $K_{\mathrm{D1}}$ and $K_{\mathrm{D}}^{\mathrm{app}}$ are experimentally determined. Given it is apparent, this dissociation constant reports broadly on the nucleotide affinity. Presented rates of association and dissociation are also apparent.

**Equation for the concentration of binary $[\alpha.\mathrm{DNA}]$ complex in equilibrium:**

In the steady state (equilibrium) the following equation is valid:

$$\frac{d[\alpha.\mathrm{DNA}]}{dt} = 0$$

If the general working model is expressed through the rate equations where rate of $[\alpha.\mathrm{DNA}]$ complex formation is equal to the rate of $[\alpha.\mathrm{DNA}]$ complex dissociation, then:

$$k_1[\alpha][\mathrm{DNA}] + k_{4*}[\alpha.\mathrm{DNA.dGTP}] = k_2[\alpha.\mathrm{DNA}] + k_{3*}[\alpha.\mathrm{DNA}][\mathrm{dGTP}]$$

$$k_1[\alpha][\mathrm{DNA}] + k_{4*}[\alpha.\mathrm{DNA.dGTP}] = [\alpha.\mathrm{DNA}](k_2 + k_{3*}[\mathrm{dGTP}])$$

$$[\alpha.\mathrm{DNA}] = \frac{k_1[\alpha][\mathrm{DNA}] + k_{4*}[\alpha.\mathrm{DNA.dGTP}]}{(k_2 + k_{3*}[\mathrm{dGTP}])}$$

In the steady state, it is also valid:

$$\frac{d[\alpha.\mathrm{DNA.dGTP}]}{dt} = 0$$

If this equation is expressed through the rate equations where rate of $[\alpha.\mathrm{DNA.dGTP}]$ complex formation is equal to the rate of $[\alpha.\mathrm{DNA.dGTP}]$ complex dissociation, then:

$$k_{3*}[\alpha.\mathrm{DNA}][\mathrm{dGTP}] = k_{4*}[\alpha.\mathrm{DNA.dGTP}]$$

$$[\alpha.\mathrm{DNA}] = \frac{k_1[\alpha][\mathrm{DNA}] + k_{4*}[\alpha.\mathrm{DNA.dGTP}]}{(k_2 + k_{3*}[\mathrm{dGTP}])}$$

$$[\alpha.\mathrm{DNA}] = \frac{k_1[\alpha][\mathrm{DNA}] + k_{3*}[\alpha.\mathrm{DNA}][\mathrm{dGTP}]}{(k_2 + k_{3*}[\mathrm{dGTP}])}$$

$$[\alpha.\mathrm{DNA}] = \frac{[\alpha][\mathrm{DNA}]}{\frac{k_2}{k_1} + \frac{k_{3*}}{k_1} \cdot [\mathrm{dGTP}]} + \frac{[\alpha.\mathrm{DNA}][\mathrm{dGTP}]}{\frac{k_2}{k_3} + \frac{k_{3*}}{k_{3*}} \cdot [\mathrm{dGTP}]}$$

$$[\alpha.\mathrm{DNA}] = \frac{[\alpha][\mathrm{DNA}]}{K_{\mathrm{D1}} + \frac{k_{3*}}{k_2}\cdot\frac{k_2}{k_1}\cdot[\mathrm{dGTP}]} + \frac{[\alpha.\mathrm{DNA}][\mathrm{dGTP}]}{\frac{k_2}{k_{3*}} + [\mathrm{dGTP}]}$$

$$[\alpha.\mathrm{DNA}] = \frac{[\alpha][\mathrm{DNA}]}{K_{\mathrm{D1}} + \frac{k_{3*}}{k_2}K_{\mathrm{D1}}[\mathrm{dGTP}]} + \frac{[\alpha.\mathrm{DNA}][\mathrm{dGTP}]}{\frac{k_2}{k_{3*}} + [\mathrm{dGTP}]}$$

If the new temporary constant F=$\frac{k_{3*}}{k_2}$ is introduced for simplification:

$$[\alpha.\mathrm{DNA}] = \frac{[\alpha][\mathrm{DNA}]}{K_{\mathrm{D1}} + \mathrm{F}\cdot K_{\mathrm{D1}}[\mathrm{dGTP}]} + \frac{[\alpha.\mathrm{DNA}][\mathrm{dGTP}]}{\frac{1}{\mathrm{F}} + [\mathrm{dGTP}]}$$

$$[\alpha.\mathrm{DNA}]\left(1 - \frac{[\mathrm{dGTP}]}{\frac{1}{\mathrm{F}} + [\mathrm{dGTP}]}\right) = \frac{[\alpha][\mathrm{DNA}]}{K_{\mathrm{D1}} + \mathrm{F}\cdot K_{\mathrm{D1}}[\mathrm{dGTP}]}$$

$$[\alpha.\mathrm{DNA}]\left(1 - \frac{[\mathrm{dGTP}]}{\frac{1 + \mathrm{F}[\mathrm{dGTP}]}{\mathrm{F}}}\right) = \frac{[\alpha][\mathrm{DNA}]}{K_{\mathrm{D1}} + \mathrm{F}\cdot K_{\mathrm{D1}}[\mathrm{dGTP}]}$$

$$[\alpha.\mathrm{DNA}]\left(1 - \frac{\mathrm{F}[\mathrm{dGTP}]}{1 + \mathrm{F}[\mathrm{dGTP}]}\right) = \frac{[\alpha][\mathrm{DNA}]}{K_{\mathrm{D1}}(1 + \mathrm{F}[\mathrm{dGTP}])}$$

$$[\alpha.\mathrm{DNA}]\left(\frac{1 + \cancel{\mathrm{F}[\mathrm{dGTP}]} - \cancel{\mathrm{F}[\mathrm{dGTP}]}}{1 + \mathrm{F}[\mathrm{dGTP}]}\right) = \frac{[\alpha][\mathrm{DNA}]}{K_{\mathrm{D1}}(1 + \mathrm{F}[\mathrm{dGTP}])}$$

$$[\alpha.\mathrm{DNA}]\left(\frac{1}{\cancel{1 + \mathrm{F}[\mathrm{dGTP}]}}\right) = \frac{[\alpha][\mathrm{DNA}]}{K_{\mathrm{D1}}(\cancel{1 + \mathrm{F}[\mathrm{dGTP}]})}$$

$$\boxed{[\alpha.\mathrm{DNA}] = \frac{[\alpha][\mathrm{DNA}]}{K_{\mathrm{D1}}}} \qquad \text{(Equation 2)}$$

**Expressing the concentration of ternary $\alpha.\mathbf{DNA}.\mathbf{dGTP}$ complex in equilibrium**

$$k_{4*}[\alpha.\mathrm{DNA}.\mathrm{dGTP}] = k_{3*}[\alpha.\mathrm{DNA}][\mathrm{dGTP}]$$

$$[\alpha.\mathrm{DNA}.\mathrm{dGTP}] = \frac{k_{3*}[\alpha.\mathrm{DNA}][\mathrm{dGTP}]}{k_{4*}} = \frac{[\alpha.\mathrm{DNA}][\mathrm{dGTP}]}{\frac{k_{4*}}{k_{3*}}} = \frac{[\alpha.\mathrm{DNA}][\mathrm{dGTP}]}{K_{\mathrm{D2}}^{\mathrm{app}}}$$

When $[\alpha.\mathrm{DNA}]$ from Equation 2 is substituted into the previous expression:

$$\boxed{[\alpha.\mathrm{DNA}.\mathrm{dGTP}] = \frac{[\alpha.\mathrm{DNA}][\mathrm{dGTP}]}{K_{\mathrm{D2}}^{\mathrm{app}}} = \frac{[\alpha][\mathrm{DNA}][\mathrm{dGTP}]}{K_{\mathrm{D1}}\cdot K_{\mathrm{D2}}^{\mathrm{app}}}} \qquad \text{(Equation 3)}$$

**Expressing the sum of concentrations of binary and ternary complexes in equilibrium on the chip surface**

In equilibrium, the two species (binary and ternary complex) may be simultaneously present bound to the immobilised p/t DNA on the chip surface, so the total mass (concentration) of bound species equates to sum of the two (note that the masses of binary and ternary complexes are essentially the same given that mass of bound dGTP is small relative to the mass of $\alpha.\mathrm{DNA}$; hence, mass and molar concentrations are inter-convertible:

$$[\alpha.\mathrm{DNA}] + [\alpha.\mathrm{DNA.dGTP}] = \frac{[\alpha][\mathrm{DNA}]}{K_{\mathrm{D1}}} + \frac{[\alpha][\mathrm{DNA}][\mathrm{dGTP}]}{K_{\mathrm{D1}} \cdot K_{\mathrm{D2}}^{\mathrm{app}}}$$

$$[\alpha.\mathrm{DNA}] + [\alpha.\mathrm{DNA.dGTP}] = \frac{[\alpha][\mathrm{DNA}]}{K_{\mathrm{D1}}}\left(1 + \frac{[\mathrm{dGTP}]}{K_{\mathrm{D2}}^{\mathrm{app}}}\right) \quad \text{(Equation 4)}$$

In equilibrium, the sum of two species on the surface corresponds to measured SPR response $R_{eq}$:

$$[\alpha.\mathrm{DNA}] + [\alpha.\mathrm{DNA.dGTP}] = R_{\mathrm{eq}} \quad \text{(Equation 5)}$$

$[\mathrm{DNA}]$ represents the concentration of free ligand immobilised on the chip surface. During interaction, some of the ligand sites become occupied with bound $\alpha$ and/or $\alpha.\mathrm{dGTP}$ so the concentration of free $[\mathrm{DNA}]$ ligand in equilibrium is $[\mathrm{DNA}] = [\mathrm{DNA}]_0 - [\alpha.\mathrm{DNA}] - [\alpha.\mathrm{DNA.dGTP}]$, where $[\mathrm{DNA}]_0$ is the initial concentration of the free ligand at the start of the reaction. When Equation 5 is substituted into the previous equation, the free ligand concentration at equilibrium is $[\mathrm{DNA}] = [\mathrm{DNA}]_0 - R_{\mathrm{eq}}$.

When all ligand sites are occupied (at saturation), the free ligand concentration on the surface must be $[\mathrm{DNA}] = 0$ while the measured response at equilibrium $R_{\mathrm{eq}}$ is maximal and denoted $R_{\mathrm{max}}$. Because of the 1:1 molar relationship between unreacted and reacted DNA (in our case just distributed between binary and ternary species on the surface), the initial concentration of ligand on the surface (at time zero) can be relatively expressed in terms of SPR response units (RU) as exactly corresponding to maximum binding capacity $R_{\mathrm{max}}$, that is $[\mathrm{DNA}]_0 = R_{\mathrm{max}}$. Using this substitution and substitution from Equation 5, it follows that the expression for unreacted DNA concentration at equilibrium $[\mathrm{DNA}] = [\mathrm{DNA}]_0 - [\alpha.\mathrm{DNA}] - [\alpha.\mathrm{DNA.dGTP}]$ can be simply rewritten as:

$$[\mathrm{DNA}] = R_{\mathrm{max}} - R_{\mathrm{eq}} \quad \text{(Equation 6)}$$

When Equation 5 and Equation 6 are substituted into Equation 4:

$$R_{\mathrm{eq}} = \frac{[\alpha]}{K_{\mathrm{D1}}}\left(R_{\mathrm{max}} - R_{\mathrm{eq}}\right)\left(1 + \frac{[\mathrm{dGTP}]}{K_{\mathrm{D2}}^{\mathrm{app}}}\right)$$

$$R_{\mathrm{eq}}\ (1 + \frac{[\alpha]}{K_{\mathrm{D1}}}\left(1 + \frac{[\mathrm{dGTP}]}{K_{\mathrm{D2}}^{\mathrm{app}}}\right) = R_{\mathrm{max}} \cdot \frac{[\alpha]}{K_{\mathrm{D1}}}\left(1 + \frac{[\mathrm{dGTP}]}{K_{\mathrm{D2}}^{\mathrm{app}}}\right)$$

$$\boxed{R_{\mathrm{eq}} = R_{\mathrm{max}}\left(\frac{\frac{[\alpha]}{K_{\mathrm{D1}}}\left(1 + \frac{[\mathrm{dGTP}]}{K_{\mathrm{D2}}^{\mathrm{app}}}\right)}{1 + \frac{[\alpha]}{K_{\mathrm{D1}}}\left(1 + \frac{[\mathrm{dGTP}]}{K_{\mathrm{D2}}^{\mathrm{app}}}\right)}\right)} \quad \text{(Equation 7)}$$

In case of a constant [dGTP], a situation we have in SPR experiments given that constant concentration ([dGTP] = 0.5 mM) is maintained in the flow, a new constant…

$$1 + \frac{[\mathrm{dGTP}]}{K_{\mathrm{D2}}^{\mathrm{app}}} = \mathrm{C}$$

…can be introduced into Equation 7 for simplification:

$$R_{\mathrm{eq}} = R_{\mathrm{max}}\left(\frac{\frac{[\alpha]}{K_{\mathrm{D1}}} \cdot \mathrm{C}}{1 + \frac{[\alpha]}{K_{\mathrm{D1}}} \cdot \mathrm{C}}\right)$$

$$R_{\mathrm{eq}} = R_{\max}\left(\frac{\frac{\mathrm{C}[\alpha]}{K_{\mathrm{D1}}}}{\frac{K_{\mathrm{D1}} + \mathrm{C}[\alpha]}{K_{\mathrm{D1}}}}\right) = R_{\max}\left(\frac{\mathrm{C}[\alpha]}{K_{\mathrm{D1}} + \mathrm{C}[\alpha]}\right) = R_{\max}\left(\frac{[\alpha]}{[\alpha] + \frac{K_{\mathrm{D1}}}{\mathrm{C}}}\right)$$

When constant C is back-substituted and a new constant $K_{\mathrm{D}}^{\mathrm{app}}$ introduced, we obtain:

$$R_{\mathrm{eq}} = R_{\max}\left(\frac{[\alpha]}{[\alpha] + \frac{K_{\mathrm{D1}}}{1 + \frac{[\mathrm{dGTP}]}{K_{\mathrm{D2}}^{\mathrm{app}}}}}\right) = R_{\max}\left(\frac{[\alpha]}{[\alpha] + K_{\mathrm{D}}^{\mathrm{app}}}\right), \quad \text{(Equation 8)}$$

where:

$$K_{\mathrm{D}}^{\mathrm{app}} = \frac{K_{\mathrm{D1}}}{1 + \frac{[\mathrm{dGTP}]}{K_{\mathrm{D2}}^{\mathrm{app}}}} \quad \text{(Equation 9)}$$

Equation 8 is of the same hyperbolic pattern as a steady-state affinity (SSA) model. Therefore measured $R_{\mathrm{eq}}$ values as a function of $[\alpha]$ concentration at fixed [dGTP] can be used to derive the $K_{\mathrm{D}}^{\mathrm{app}}$ value for the $\alpha - \mathrm{DNA}$ interaction using a simple SSA model.

Also, once $K_{\mathrm{D}}^{\mathrm{app}}$ is determined and $K_{\mathrm{D1}}$ is known, *i.e.* from its determination from the interaction between α and DNA in the absence of dGTP, rearranged Equation 9 can be used to calculate the apparent dissociation constant $K_{\mathrm{D2}}^{\mathrm{app}}$ for the interaction between α.DNA and dGTP, which is thus globally related to the nucleotide affinity:

$$K_{\mathrm{D2}}^{\mathrm{app}} = \frac{K_{\mathrm{D}}^{\mathrm{app}}[\mathrm{dGTP}]}{K_{\mathrm{D1}} - K_{\mathrm{D}}^{\mathrm{app}}} \quad \text{(Equation 10)}$$

**Supplementary Data 2: Summary and extended discussions of SPR studies on α binding to p/t DNAs**

| Reaction | ss DNA | p/t DNA 0–2 nt | p/t DNA 0–4 nt | p/t DNA 0–5 nt | p/t DNA 0–6 nt | p/t DNA 0–7 nt | p/t DNA 0–9 nt | p/t DNA 0–14 nt |
|---|---|---|---|---|---|---|---|---|
| **$K_{\mathrm{D1}}$ (nM)** $\alpha^{\mathrm{E612K}}$–DNA | 1200 ± 130 | 3800 ± 400 | 1800 ± 200 | 1300 ± 100 | 630 ± 50 | 430 ± 30 | 280 ± 30 | 240 ± 20 |
| **$K_{\mathrm{D}}^{\mathrm{app}}$ (nM)** $\alpha^{\mathrm{E612K}}$–DNA + dGTP | 890 ± 80 | 3500 ± 400 | 890 ± 50 | 370 ± 20 | 68 ± 6 | 43 ± 4 | 20 ± 4 | 14 ± 2 |
| **Calculated $K_{\mathrm{D2}}^{\mathrm{app}}$ (μM)** $\alpha^{\mathrm{E612K}}$.DNA–dGTP | – | > 1535 | 500 ± 200 | 190 ± 50 | 61 ± 12 | 56 ± 10 | 42 ± 14 | 32 ± 8 |
| **$K_{\mathrm{D1}}$ (nM)** wt α–DNA | – | – | – | – | – | – | 3800 ± 400 | 3100 ± 200 |
| **$K_{\mathrm{D}}^{\mathrm{app}}$ (nM)** wt α–DNA | – | – | – | – | – | – | 1110 ± 60 | 820 ± 50 |
| **Calculated $K_{\mathrm{D2}}^{\mathrm{app}}$ (μM)** wt α.DNA–dGTP | – | – | – | – | – | – | 210 ± 50 | 180 ± 30 |

**Table 1. Equilibrium binding parameters of $\alpha^{E612K}$ and wild-type $\alpha$ binding to p/t DNAs with ss template overhang of various lengths in the absence and presence of dGTP, based on SPR data in Supplementary Figures 2, 3 and 4.** Dissociation constants for the $^{bio}$p/t DNA−$\alpha$ interaction ($K_{D1}$) and apparent dissociation constant ($K_D^{app}$**)** for the $^{bio}$p/t DNA−$\alpha$ interactions (in the presence of 0.5 mM dGTP) were determined by fitting responses at equilibrium in Supplementary Figures 2, 3 and 4 to Equation 1 (Supplementary Methods) and Equation 8 (Supplementary Data 1), respectively. The apparent dissociation constant $K_{D2}^{app}$ that broadly reports on the nucleotide affinity was calculated from Equation 10 (Supplementary Data 1). The errors are standard errors of the fits. 0–2 nt contains a total of 3 nt template overhang in the absence of dGTP, but 2 nt in presence of dGTP, whereby 0 nt stands for the templating nucleotide that pairs with incoming nucleotide.

Data in Table 1 reveal the following:

1) At "saturating" ssDNA lengths (p/t DNAs with 10 or more nt on the ss overhang), comparison of $K_{D1}$ values shows that $\alpha^{E612K}$ (240 ± 20 nM) binds DNA up to 15-fold stronger than wt $\alpha$ (3100 ± 200 nM).

2) Comparison of $K_{D1}$ values shows similar stabilities between $\alpha^{E612K}$–DNA (0–2 nt) and wt $\alpha$–DNA (0–9 nt) binary complexes (both 3800 ± 400 nM), suggesting that interaction of residue $\alpha$612K with −2 nt of p/t DNA (Figure 2D) significantly stabilizes the binary complex. Such stabilization, that also extends to the ternary complex (this work), can partially compensate for the general need for stabilisation of the ternary complex by the $\beta_2$-clamp in replication assays, i.e. $\alpha^{E612K}$ was able to replicate large stretches of DNA processively even in the absence of $\beta_2$, while wt $\alpha$ was not (Yanagihara et all, 2007).

3) Relative to previous point 2, we note that in the presence of dGTP, $K_D^{app}$ value for $\alpha^{E612K}$–DNA (0–2 nt; 3500 ± 400 nM) remained largely unperturbed relative to its $K_{D1}$ (3800 ± 400 nM), likely suggesting either failure to produce practically any ternary complex at 0.5 mM dGTP. In contrast $K_D^{app}$ for wt $\alpha$–DNA (0–9 nt; 1110 ± 60 nM) reduced almost 4-fold relative to $K_{D1}$ (3800 ± 400 nM). Given their $K_{D1}$ values are similar while wt $\alpha$ is a less potent polymerase compared to $\alpha^{E612K}$, these results demonstrate the critical importance that contacts between a sufficiently long ss region of DNA template and polymerase play (structurally and kinetically) in converting $\alpha$.DNA to the productive ternary state (transit to polymerization mode).

4) Measured $K_D^{app}$ values for $\alpha^{E612K}$–DNA in the presence of dGTP reveal a relatively minor contribution to the stability of the ternary complex when more than six residues ahead of the templating nucleotide are utilized in the interaction. Nevertheless, it seems that ~10 ss residues can still be sporadically engaged.

Next, the data indicate a strong ~13-fold reduction of $K_D^{app}$ value in $\alpha^{E612K}$–DNA interaction (from 890 ± 50 nM to 68 ± 6 nM) when the overhang extended from 4 to 6 nt ahead of templating base, which could in the first instance indicate strong preference of $\alpha^{TC}$ to stably engage not only first 4 nt but also 5$^{th}$ and 6$^{th}$ nt. Yet, in the structures of PolIII$^{TC}$ and molecular dynamics simulation of ternary state only 4 nt ahead of templating base were typically seen in stable contact with $\alpha$ reaching OB.

We note that in both PolIII$^{TC}$ structures and molecular dynamics simulation, [dGTP] was at saturation – in structures because of presence of other stabilising interactions connected to DNA-loaded $\beta_2$ as well as the sufficient overhang length (50 nt) relative to utilized 1 mM [dGTP] (as SPR results would indicate), and in simulation because of the direct placement of dGTP in the nucleotide binding pocket. However, $K_{D2}^{app}$ value for overhang extending 4 nt ahead of templating nucleotide (500 ± 20 $\mu$M) indicates that utilised [dGTP] of 500 $\mu$M is far from saturating.

In order to properly compare true contribution of each additional overhang nucleotide from 4th to 6th nt to the stability of the ternary $\alpha^{E612K}$•DNA•dGTP complex, we decided to even out the comparison landscape by cancelling out the effect of variability in dGTP saturation levels. To do that, we opted to normalise relevant $K_D^{app}$ values to the dGTP saturation level equivalent to that of the template overhang with 6 nt ahead of templating nucleotide (68 ± 6 nM). We first calculated the nucleotide saturation level factor $\frac{[\mathrm{dGTP}]}{K_{\mathrm{D2}}^{\mathrm{app}}}$ (from Equation 9) for this overhang:

$$\frac{[\mathrm{dGTP}]}{K_{\mathrm{D2}}^{\mathrm{app}}} = \frac{500\ \mu\mathrm{M}}{61 \pm 12\ \mu\mathrm{M}} = 8 \pm 2,$$

then applied the same factor and Equation 9 to normalise $K_D^{app}$ value for $\alpha^{E612K}$–DNA interaction with 4 nt overhang:

$$K_{\mathrm{D}}^{\mathrm{app,N4}} = \frac{K_{\mathrm{D1}}}{1 + \frac{[\mathrm{dGTP}]}{K_{\mathrm{D2}}^{\mathrm{app}}}} = \frac{1800 \pm 200\ \mathrm{nM}}{1 + (8 \pm 2)} = \frac{1800 \pm 200\ \mathrm{nM}}{9 \pm 2} = 200 \pm 70\ \mathrm{nM},$$

and with 5 nt overhang ($K_D^{app,N5}$ = 140 ± 40 nM). Taking errors into account, the minimal preference for $\alpha$ (OB) to engage both 5th and 6th overhang nucleotide beyond first 4 appears relatively modest < 2-fold ($\frac{K_{\mathrm{D}}^{\mathrm{app,N4}}}{K_{\mathrm{D}}^{\mathrm{app,6}}} = \frac{200 - 70\ \mathrm{nM}}{68+6\ \mathrm{nM}} = 1.8$), with also modest minimal preference (1.4-fold) to engage 5th nucleotide. Lack of much stronger preference to engage these overhang nucleotides in interaction with $\alpha$ could explain lack of their detection (stable interaction) in PolIII$^{TC}$ (Figure 3D), or their rather more sporadic engagement in interaction with $\alpha$ as their constitutive overhang segment wiggles between various bound and unbound states (Figure 5B–D). Taken together, our structural, molecular dynamics simulation and SPR studies indicate that physiologically (most) relevant contact of $\alpha$ with ss overhang of p/t DNA likely involves 4 nt ahead of templating nucleotide.

Notably, $\alpha$ (OB) still shows a 4.4-fold maximal preference ($\frac{K_{\mathrm{D}}^{\mathrm{app,N4}}}{K_{\mathrm{D}}^{\mathrm{app,6}}} = \frac{200 + 70\ \mathrm{nM}}{68 - 6\ \mathrm{nM}} = 4.4$) for engaging the 5th and 6th overhang nucleotides compared to the first 4. It could be that these residues more actively participate in binding when overhang lengths are restricted. In these cases, reduced degrees of freedom compared to longer overhangs could prevent these nucleotides from delocalization (as seen in molecular dynamic simulation), which could stabilise their interactions with OB. Consequently, $\alpha$ may as well engage the 5th and 6th overhang nucleotides ahead of templating base *in vivo*.
